# Brain-age prediction: a systematic comparison of machine learning workflows

**DOI:** 10.1101/2022.11.16.515405

**Authors:** Shammi More, Georgios Antonopoulos, Felix Hoffstaedter, Julian Caspers, Simon B. Eickhoff, Kaustubh R. Patil, the Alzheimer’s Disease Neuroimaging Initiative

## Abstract

The difference between age predicted using anatomical brain scans and chronological age, i.e., the brain-age delta, provides a proxy for atypical aging. Various data representations and machine learning (ML) algorithms have been used for brain-age estimation. However, how these choices compare on performance criteria important for real-world applications, such as; (1) within-site accuracy, (2) cross-site generalization, (3) test-retest reliability, and (4) longitudinal consistency, remains uncharacterized. We evaluated 128 workflows consisting of 16 feature representations derived from gray matter (GM) images and eight ML algorithms with diverse inductive biases. Using four large neuroimaging databases covering the adult lifespan (total N = 2953, 18-88 years), we followed a systematic model selection procedure by sequentially applying stringent criteria. The 128 workflows showed a within-site mean absolute error (MAE) between 4.73-8.38 years, from which 32 broadly sampled workflows showed a cross-site MAE between 5.23-8.98 years. The test-retest reliability and longitudinal consistency of the top 10 workflows were comparable. The choice of feature representation and the ML algorithm both affected the performance. Specifically, voxel-wise feature spaces (smoothed and resampled), with and without principal components analysis, with non-linear and kernel-based ML algorithms performed well. Strikingly, the correlation of brain-age delta with behavioral measures disagreed between within-site and cross-site predictions. Application of the best-performing workflow on the ADNI sample showed a significantly higher brain-age delta in Alzheimer’s and mild cognitive impairment patients. However, in the presence of age bias, the delta estimates in the diseased population varied depending on the sample used for bias correction. Taken together, brain-age shows promise, but further evaluation and improvements are needed for its real-world application.

**Highlights:**

- There is an effect of both feature space and ML algorithm on prediction error.
- Voxel-wise features performed better than parcel-wise features.
- GPR, KRR and RVR algorithms performed well.
- The within-site and cross-site delta-behavior correlations disagree.
- Higher brain-age delta inference in AD depends on data used for bias correction.

## 1. Introduction

Precision and preventive medicine, e.g., early detection of Alzheimer’s disease (AD), can benefit from individual-level quantification of atypical aging. Machine learning (ML) approaches, together with large neuroimaging datasets can provide such individualized predictions. Indeed, ML algorithms can capture the multivariate pattern of age-related changes in the brain associated with healthy or typical aging (Franke *et al*., 2010; Varikuti *et al*., 2018; Cole, 2020; Beheshti *et al*., 2022; Hahn *et al*., 2022). Such a model can then be used to predict age, i.e., brain-age, from an unseen subject’s image. Being a normative model, a large deviation between the chronological and the predicted age is indicative of atypical aging. A higher positive difference between the brain-age and chronological age, i.e., brain-age delta (which we refer to simply as delta), indicates “older-appearing” brains. As an indicator of future risk of experiencing age-associated health issues (Gaser *et al*., 2013; Cole and Franke, 2017; Richard *et al*., 2018), delta quantitatively relates to several age-related risk factors and general physical health, such as weaker grip strength, poorer lung function, history of stroke, greater frequency of alcohol intake, increased mortality risk (Cole *et al*., 2018; Cole, 2020), and poorer cognitive functions such as fluid intelligence, processing speed, semantic verbal fluency, visual attention, and cognitive flexibility (Cole *et al*., 2018; Richard *et al*., 2018; Boyle *et al*., 2021). Overall, the delta can serve as an omnibus biomarker of brain integrity and health.

Studies have shown global and local gray matter volume (GMV) loss (Good *et al*., 2001; Galluzzi *et al*., 2008; Giorgio *et al*., 2010) with aging and accelerated loss in neurodegenerative disorders (Good *et al*., 2001; Karas *et al*., 2004; Fjell *et al*., 2014). This makes GMV a clinically relevant candidate for the investigation of atypical aging via brain-age estimation (Franke *et al*., 2010; Cole *et al*., 2015). Studies using voxel-based morphometry (VBM)-derived GMV to predict brain-age have claimed prediction errors of ∼5-8 years in healthy individuals (Table S1). However, it is difficult to compare these studies as they differ in experimental setup and methodology, such as feature space used, ML algorithms, age range, and evaluation criteria. For a brain-age estimation model to be used in real-world applications, it must perform well on several evaluation criteria; (1) in addition to performing well on new data from training site, a model should perform well on data from novel sites, (2) brain-age estimation must be reliable, i.e., it should make consistent predictions on repeated measurements, and (3) it should also exhibit longitudinal consistency, i.e., the predicted age should be proportionally higher for later scans after a longer duration.

A brain-age estimation workflow consists of a feature space and an ML algorithm, and several choices exist for each. One can choose voxel-wise data with additional smoothing and/or resampling. Additionally, dimensionality reduction methods such as principal components analysis (PCA) can be applied, which can improve the observations-to-features ratio and signal-to-noise ratio (Franke *et al*., 2010, 2013; Gaser *et al*., 2013). Another option to reduce dimensionality is averaging voxels in each parcel within a brain atlas (Varikuti *et al*., 2018; Eickhoff *et al*., 2021). One also needs to choose from a large pool of ML algorithms, such as random forest regression (RFR), relevance vector regression (RVR), and Gaussian process regression (GPR), many of which have shown success in brain-age estimation. These choices are known to affect performance (Gutierrez Becker *et al*., 2018, Baecker *et al*., 2021*a*; de Lange *et al*., 2022). However, previous studies have performed only limited comparisons on the same data and setup.

A critical aspect, especially for clinical application, is the commonly reported negative correlation between delta and true age (Beheshti *et al*., 2019, Smith *et al*., 2019*a*; de Lange and Cole, 2020). This may result in spurious correlations between the delta and non-imaging measures, making it difficult to interpret relationships when chronological age is not accounted for (Franke *et al*., 2013; Löwe *et al*., 2016). This age bias complicates or may even mislead downstream individualized decision-making. It can be mitigated using bias correction models; usually, linear regression predicting brain-age or delta using chronological age (Le *et al*., 2018; Liang *et al*., 2019, Smith *et al*., 2019*b*; de Lange *et al*., 2022). These correction models are also used to counter the systematic under- or over-estimation of age in a novel site, usually reflected in the non-zero average delta in healthy controls. Either data from the training samples or the application site, especially the healthy controls, are used to obtain bias correction models. Furthermore, the sample size has substantial impact on quality of the model. However, the impact of these choices have not been fully investigated. Taken together, there is a lack of understanding regarding the choices for designing a brain-age estimation workflow and the estimation of individual-level delta.

To fill this gap, we systematically assessed 128 workflows consisting of 16 feature spaces derived from gray matter (GM) images and eight ML algorithms with diverse inductive biases to establish guidelines for brain-age estimation workflow design. Using several large neuroimaging databases with a wide age range coverage, we first evaluated these workflows for their within-site (or single-site) and cross-site performances. Next, we evaluated the test-retest reliability and longitudinal consistency of some top-performing workflows. Then, we assessed the performance of our best-performing workflow in a clinical sample. We investigated the correlations between delta and behavioral/cognitive measures in healthy and clinical cohorts and various factors affecting these correlations. We also compared our best-performing workflow with a publicly available model, brainageR, which is trained on three tissue types, GM, white matter (WM), and Cerebrospinal fluid (CSF), with the Gaussian processes regression (GPR) algorithm. Several follow-up analyses were performed to investigate the effect of preprocessing (CAT vs. SPM) and tissue type (GM vs. GM+WM+CSF) choices on prediction performance.

## 2. Material and Methods

### 2.1 Datasets

#### 2.1.1 MRI data

We used T1-weighted (T1w) magnetic resonance imaging (MRI) data from healthy subjects covering a wide age range (18-88 years, training data) from several large neuroimaging datasets (Table 1), including the Cambridge Centre for Ageing and Neuroscience (CamCAN, N = 651) (Taylor *et al*., 2017), Information eXtraction from Images (IXI, N = 562) (https://brain-development.org/ixi-dataset/), the enhanced Nathan Kline Institute-Rockland Sample (eNKI, N = 597) (Nooner *et al*., 2012), the 1000 brains study (1000BRAINS; N = 1143) (Caspers *et al*., 2014), Consortium for Reliability and Reproducibility (CoRR) (Zuo *et al*., 2014), the Open Access Series of Imaging Studies (OASIS-3) (LaMontagne *et al*., 2019), and the MyConnectome dataset (Poldrack *et al*., 2015). The inclusion criteria were age between 18 and 90 years, gender data available, and no current or past known diagnosis of neurological, psychiatric, or major medical conditions. The IXI dataset was acquired from multiple sites; however, we treat it as a single dataset representing typical data acquired in a noisy clinical setting. From the OASIS-3 dataset, we selected scans from cognitively normal subjects acquired on 3T scanners.

**Table 1.** Sample characteristics of the datasets used in the current study. Datasets used a. for training single-site or within-site models. b. for training cross-site models. c. to evaluate test-retest reliability and longitudinal consistency of brain-age delta and comparison with brainageR. d. to evaluate performance in clinical samples. Abbreviations: CamCAN: the Cambridge Centre for Ageing and Neuroscience, IXI: Information eXtraction from Images (includes 1.5 and 3T scans), eNKI: the enhanced Nathan Kline Institute-Rockland Sample, CoRR: Consortium for Reliability and Reproducibility, OASIS-3: the Open Access Series of Imaging Studies, ADNI: the Alzheimer’s Disease Neuroimaging Initiative, CN: cognitively normal, EMCI and LMCI: early and late mild cognitively impaired, AD: Alzheimer’s disease

| Train dataset | No. of subjects (N) | Males/Females | Age range | Mean ± S.D. | Median |
| --- | --- | --- | --- | --- | --- |
| CamCAN | 651 | 321/330 | 18 - 88 | 54.27 ± 18.58 | 54.50 |
| IXI | 562 | 249/313 | 20 - 86 | 48.70 ± 16.44 | 48.85 |
| eNKI | 597 | 188/409 | 18 - 85 | 48.25 ± 18.51 | 50.00 |
| 1000BRAINS | 1143 | 660/513 | 22 - 85 | 61.85 ± 12.39 | 63.60 |

b.
| Train dataset | Train N | Test dataset | Test N |
| --- | --- | --- | --- |
| IXI + eNKI + 1000BRAINS | 2302 | CamCAN | 651 |
| CamCAN + eNKI + 1000BRAINS | 2391 | IXI | 562 |
| IXI + CamCAN + 1000BRAINS | 2356 | eNKI | 597 |
| IXI + CamCAN + eNKI | 1810 | 1000BRAINS | 1143 |
| IXI + CamCAN + eNKI + 1000BRAINS | 2953 | CoRR, OASIS-3, MyConnectome, ADNI | See below (c & d) |

c.
| Dataset | Data Filtering | N | Males/<br>Females | Age Range | Mean ± S.D. | Median |
| --- | --- | --- | --- | --- | --- | --- |
| CoRR | Retest < 3 months | 86 | 39/47 | 20.0 - 84.0 | 48.82 ± 18.28 | 49.00 |
| | Retest 1 – 2 years | 95 | 52/43 | 18.0 - 88.0 | 34.43 ± 22.51 | 20.00 |
| | Retest 2 – 3.25 years | 26 | 18/8 | 18.0 - 57.0 | 28.09 ± 11.89 | 24.50 |
| OASIS-3 | Retest < 3 months | 36 | 21/15 | 42.66 - 80.90 | 63.46 ± 8.80 | 62.93 |
| | Retest 3- 4 years | 127 | 52/75 | 46.04 - 86.21 | 65.59 ± 8.39 | 65.90 |
| | Full sample | 806 | 338/468 | 43.00 - 89.00 | 69.07 ± 9.06 | 69.00 |
| MyConnectome | Retest < 3 years | 20 | 20/0 | 45.39 - 48.02 | 45.73 ± 0.58 | 45.56 |

d.
| Dataset | Disease | N | Males/<br>Females | Age Range | Mean ± S.D. | Median |
| --- | --- | --- | --- | --- | --- | --- |
| ADNI<br>(Timepoint-1) | CN | 209 | 99/110 | 56.3 - 94.7 | 75.67 ± 6.94 | 75.50 |
| | EMCI | 237 | 128/109 | 55.7 - 88.7 | 70.88 ± 7.12 | 70.40 |
| | LMCI | 128 | 62/65 | 55.1 - 91.5 | 72.02 ± 7.89 | 72.55 |
| | AD | 125 | 65/60 | 56.0 - 91.0 | 74.68 ± 7.99 | 75.40 |
| ADNI<br>(Timepoint-2) | CN | 153 | 70/83 | 57.3 - 95.8 | 75.89 ± 6.63 | 75.50 |
| | EMCI | 197 | 108/89 | 56.7 - 90.4 | 71.81 ± 7.04 | 71.10 |
| | LMCI | 104 | 51/53 | 56.1 - 92.5 | 73.36 ± 7.92 | 73.95 |
| | AD | 61 | 32/29 | 57.0 - 93.0 | 75.79 ± 7.83 | 76.80 |

Furthermore, we included the Alzheimer’s Disease Neuroimaging Initiative (ADNI; https://adni.loni.usc.edu/) database to evaluate the utility of brain-age in neurodegenerative disorders (Jack *et al*., 2008; Petersen *et al*., 2010). We included 3T T1w images from cognitively normal subjects (CN, N = 209), early and late mild cognitively impaired subjects (EMCI, N = 237; LMCI, N = 128), and Alzheimer’s disease (AD, N = 125) subjects. For some of these subjects, data were available for the second timepoint 1-2 years apart (CN, N = 153; EMCI, N = 197; LMCI, N = 104; AD, N = 61) (Table 1d).

#### 2.1.2 Non-imaging data

We used various behavioral/cognitive measures available in some datasets to compute their correlations with delta. The CamCAN dataset provides a measure for fluid intelligence (FI; N = 631) assessed by the Cattell Culture Fair test and reaction time for the motor learning task (N = 302) (Taylor *et al*., 2017). The eNKI dataset provides various measures for cognitive and executive functioning assessed by the Delis-Kaplan Executive Functioning System (D-KEFS) and cognitive intelligence measured by Wechsler Abbreviated Scale of Intelligence (WASI-II) (Nooner *et al*., 2012). Specifically, we used a. the Color-Word Interference Test (CWIT) inhibition trial completion time (N = 340), b. the Trail Making Test (TMT) number-letter switching condition completion time (N = 344), c. WASI-II matrix reasoning scores (N = 347), and d. WASI-II similarities scores (N = 347).

ADNI provides cognitive tests measuring disease severity such as Mini-Mental State Examination (MMSE; with lower scores being related to higher disease severity), Global Clinical Dementia Rating Scale (CDR; range 0–3, with 0 denoting CN, 0.5 denoting MCI, and a score of 1 or above denoting AD), and Functional Assessment Questionnaire (FAQ; range 0-30, with 0 scored as ‘no impairment’ and 30 as ‘severely impaired’).

All the datasets except the 1000BRAINS data are available publicly. Ethical approval and informed consent were obtained locally for each study covering both participation and subsequent data sharing. The ethics proposals for the use and retrospective analyses of the datasets were approved by the Ethics Committee of the Medical Faculty at the Heinrich-Heine-University Düsseldorf.

### 2.2 Data preparation

All T1w images were preprocessed using the VBM implementation of the Computational Anatomy Toolbox (CAT) version 12.8 (Gaser *et al*., 2022). To ensure accurate normalization and segmentation, initial affine registration of T1w images was done with higher than default accuracy (accstr = 0.8). After bias field correction and tissue class segmentation, accurate optimized Geodesic shooting (Ashburner and Friston, 2011) was used for normalization (regstr = 1). We used 1 mm Geodesic Shooting templates and outputted 1 mm isotropic images instead of the default 1.5 mm. To derive voxel-wise GMV, the normalized GM segments were modulated for linear and non-linear transformations.

For comparison with brainageR, we used corresponding preprocessing as implemented using SPM12 in MATLAB R2017b, which outputs three tissue segmentations (GM, WM, and CSF; see https://github.com/james-cole/brainageR/).

### 2.3 Workflows

Each workflow consists of a feature representation and an ML algorithm. We evaluated 128 workflows constituting 16 feature representations and eight ML algorithms.

#### 2.3.1 Feature representations

The 16 feature representations were derived from the CAT-preprocessed voxel-wise GM images. Using voxel-wise data directly can be problematic as it can lead to overfitting due to the curse of dimensionality owing to a large number of features compared to the number of data points. Hence, we implemented two dimensionality reduction approaches.

First, a commonly used strategy is to process voxel-wise GMV by smoothing and then resampling (Franke *et al*., 2010), which may also improve the signal-to-noise ratio. Second, we used an atlas-based strategy on voxel-wise GMV to summarize data from distinct brain regions (called parcels). In total, we created 16 feature representations.

1. SX_RY: The modulated GMV was masked using a whole-brain mask, resulting in 238955 non-zero voxels. Then, smoothing (S) with an X mm FWHM Gaussian kernel and resampling (R) using linear interpolation to Y mm spatial resolution were applied to these images with X = {0, 4, 8} and Y = {4, 8}. These combinations created six feature spaces in total (S0_R4, S0_R8, S4_R4, S4_R8, S8_R4, S8_R8, SX_R4: 29852 voxels and SX_R8: 3747 voxels).
2. SX_RY + PCA: Additionally, PCA (Jolliffe, 2002) was applied to each SX_RY feature space while retaining 100% variance, creating another six representations (S0_R4 + PCA, S0_R8 + PCA, S4_R4 + PCA, S4_R8 + PCA, S8_R4 + PCA, S8_R8 + PCA).
3. Parcel-wise: Four parcel-wise feature spaces were created by combining cortical {100, 400, 800, 1200} parcels (Schaefer et al., 2018) with 36 subcortical (Fan et al., 2016) and 37 cerebellum (Buckner et al., 2011) parcels. We calculated the mean GMV of all the voxels within each parcel (173, 473, 873, and 1273 features).

#### 2.3.2 Machine learning algorithms

We included eight ML algorithms covering diverse inductive biases: ridge regression (RR), least absolute shrinkage and selection operator (LASSO) regression (LR), elastic net regression (ENR), kernel ridge regression (KRR), GPR, RFR, RVR with the linear kernel (RVRlin), and polynomial kernel of degree 1 (RVRpoly). These algorithms have been widely used in the prediction of age and other behavior variables from neuroimaging data (Franke *et al*., 2010; Gaser *et al*., 2013; Su *et al*., 2013; Cole *et al*., 2015, 2018; Varikuti *et al*., 2018; Jonsson *et al*., 2019; Liang *et al*., 2019; Zhao *et al*., 2019; Cole, 2020; He *et al*., 2020, Baecker *et al*., 2021*b*; Boyle *et al*., 2021; Lee *et al*., 2021; Peng *et al*., 2021; Treder *et al*., 2021; Vidal-Pineiro *et al*., 2021; Beheshti *et al*., 2022) (Table S1). Details of these algorithms are provided in the Supplementary Methods.

Recently, deep-learning (DL) models have been applied for brain-age estimation with success (Jiang *et al*., 2019; Jonsson *et al*., 2019; Peng *et al*., 2021). However, in this work, we focus on conventional ML models for the following reasons: (1) ML models have shown competitive performance to DL models (Cole *et al*., 2017; He *et al*., 2020; Schulz *et al*., 2020; Grinsztajn *et al*., 2022), and (2) the resources required for ML are more readily available and thus still enjoy wider applicability with a lower computational footprint (Thompson *et al*., 2020; van Wynsberghe, 2021).

#### 2.3.3 Learning setup and software

The ML algorithm’s hyperparameters were estimated in a nested fashion using an inner cross-validation (CV) (Varoquaux *et al*., 2017). Before training, features with low variance were removed (threshold < 1e-5), and the remaining features were Z-scored to have zero mean and unit variance. When PCA was applied, we retained components to explain 100% of the variance. These preprocessing steps were applied in a CV-consistent fashion to avoid data leakage, i.e., the parameters were estimated on the training set and applied to both the training and the test set (More *et al*., 2021).

All the workflows were implemented in Python version 3.9.1 using the Julearn machine-learning library (https://juaml.github.io/julearn/), which in turn uses the scikit-learn library for the learning algorithms KRR, GPR, and RFR (http://scikit-learn.org/) (Pedregosa *et al*., 2011). LR, RR, and ENR were implemented using the Python wrapper for glmnet (https://pypi.org/project/glmnet/) (Friedman *et al*., 2010). RVRlin and RVRpoly were implemented using the scikit-rvm package (https://github.com/JamesRitchie/scikit-rvm/). The codes used for preprocessing, feature extraction and model training are available at https://github.com/juaml/brainage_estimation.

### 2.4 Analysis setup

It is crucial to evaluate the performance of brain-age estimation workflows in different evaluation scenarios, as models that work well in one might be suboptimal when evaluated in another. To accommodate different real-world scenarios, we followed a systematic procedure where the workflows were subjected to increasingly stringent evaluations (Figure 1). In brief, we first evaluated the within-site CV performance of the 128 workflows. Next, 32 diverse workflows out of 128 were selected for cross-site evaluation by sampling over the complete CV performance range to allow for the possibility that workflows with low within-site performance might perform well in cross-site evaluation. Finally, the top 10 workflows out of the 32 were evaluated for their test-retest reliability and longitudinal consistency. After considering all the evaluation criteria, the best-performing workflow was chosen and used for application on ADNI data and comparison with brainageR. Specific analysis steps are described below.

**Figure 1.**
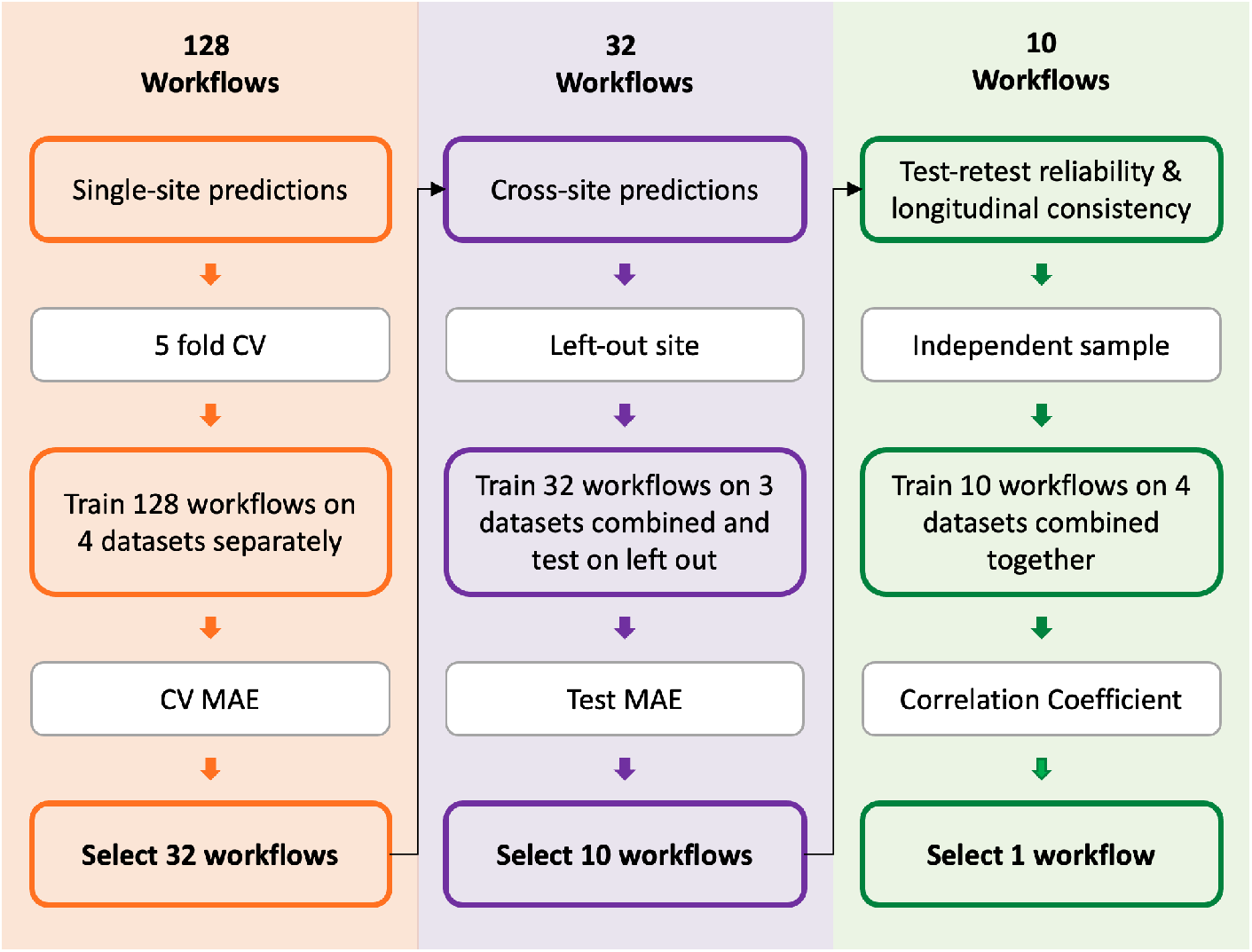
The framework to select the best-performing workflow for brain-age prediction. A total of 128 workflows were first evaluated for their single-site or within-site prediction performance using five-fold cross-validation (CV). Next, 32 workflows were selected based on the CV mean absolute error (MAE) and assessed for cross-site prediction performance. Within-site and cross-site evaluations were performed using four datasets (CamCAN, IXI, eNKI and 1000BRAINS). Then, 10 workflows out of 32 were selected based on their test MAE and assessed for test-retest reliability and longitudinal consistency using OASIS-3 and CoRR datasets. The best-performing workflow was selected after considering all the evaluation criteria.

#### 2.4.1 Within-site evaluation

We evaluated the 128 workflows (see section 2.3) separately on four datasets, CamCAN, IXI, eNKI, and 1000BRAINS. This scenario assumes that enough within-site training data are available and is widely used in brain-age estimation work (Ashburner, 2007; Su *et al*., 2013; Gutierrez Becker *et al*., 2018). To estimate a single out-of-sample brain-age for each subject, we used a 5-fold CV. For each hold-out (test) fold, the remaining 80% of the data were used for training. It is important to obtain a single prediction per subject (as opposed to multiple predictions per subject if the outer CV were repeated) for further meaningful analyses, such as correlation with non-imaging measures. Consequently, we computed two performance measures, test, and CV performance. The test performance was obtained by averaging over the 5 folds. Within each CV fold, the 80% training data were also used to obtain a generalization estimate using 5 times repeated 5-fold (5×5-fold) nested CV. All CV analysis was stratified by age to preserve the age distribution. The CV performance was obtained by first averaging over the inner 5×5-fold CV and then over the outer 5-fold CV for each dataset. Finally, the CV and test performance were averaged over the four datasets. The performance was evaluated using mean absolute error (MAE), Pearson’s correlation between predicted and true (chronological) age, and the coefficient of determination R^2^.

We followed a systematic procedure to select a subset of workflows with high accuracy while maintaining diversity. Specifically, the workflows were arranged in the increasing order of their average CV MAE and divided into 16 groups. Next, the top two workflows (with the lowest CV MAE) from each group were selected, resulting in 32 workflows. This procedure considers the possibility that workflows that are suboptimal in within-site evaluation could perform well in cross-site evaluation.

#### 2.4.2 Cross-site evaluation

Next, we tested the 32 selected workflows on cross-site data to obtain sample-unbiased performance. This emulates the real-world scenario where data from the application site are not available, and the training and test data come from different sources with confounding effects, such as scanner hardware or operator inconsistencies (Jovicich *et al*., 2006; Chen *et al*., 2014). Three out of four datasets (CamCAN, IXI, eNKI and 1000BRAINS) were pooled to form the training data, and the hold-out dataset (site) was used as the test data. A 5×5-fold CV was performed on the training data to estimate the generalization performance with an internal CV for hyperparameter tuning. The CV performance was averaged over 5×5-fold CV and then over the four hold-out sites. The test performance was averaged over the four sites. The performance was again evaluated using MAE, Pearson’s correlation between predicted and true age, and the coefficient of determination R^2^.

The 32 workflows were arranged in increasing order of their average test MAE, i.e., their average performance on the hold-out sites, from which the top 10 workflows were selected.

#### 2.4.3 Test-retest reliability and longitudinal consistency

We then trained models using the 10 selected workflows with the four datasets combined as training data (IXI + eNKI + CamCAN + 1000BRAINS, N = 2953; Supplementary Figure S1). The test-retest reliability and longitudinal consistency of the delta were evaluated for the 10 models using the OASIS-3 and CoRR datasets.

To evaluate test-retest reliability, we used: two scans from the same subjects acquired within a delay of (1) less than three months (CoRR: N = 86, age range = 20-84 years, OASIS-3: N = 36, age range = 43-81), and (2) between 1-2 years (CoRR: N = 95, age range = 18-88). The delta was calculated by subtracting chronological age (at scan time) from predicted age, and the concordance correlation coefficient (CCC) (Lin, 1989) between the delta from the two scans was calculated.

To evaluate longitudinal consistency, two scans from the same subjects acquired with a retest duration (1) between 2-3.25 years (CoRR: N = 26, age range = 18-57), and (2) between 3-4 years (OASIS-3: N = 127, age range = 46-86) were used. We computed Pearson’s correlation between the difference in the predicted age and the difference in chronological age from the two scans. A higher positive correlation here would indicate higher longitudinal consistency.

By considering the results from the within-site analysis, cross-site analysis, test-retest reliability, and longitudinal consistency, we chose one best-performing workflow for further analysis.

### 2.5 Bias Correction

Many studies have reported age-dependency of the delta with over-prediction in young subjects and under-prediction in older subjects (Le *et al*., 2018; Liang *et al*., 2019), which renders the usage of delta as an individualized biomarker problematic. A common practice is to apply a statistical bias correction to remove the effect of age from either the predicted age or the delta (Le *et al*., 2018; de Lange *et al*., 2019, Smith *et al*., 2019*b*; Cole, 2020). Note that when calculating correlations of delta with non-imaging measures, bias correction is expected to be similar to partial correlation analysis when age is used as a covariate.

Several alternatives are available for bias correction (de Lange *et al*., 2019; Liang *et al*., 2019, Smith *et al*., 2019*a*; Cole, 2020). We chose the method used by Cole and colleagues (Cole, 2020) as it does not use the chronological age of the test data, making it more meaningful for a comparative study (de Lange *et al*., 2022). Furthermore, this method is relevant for possible future applications like forensic investigations. A linear regression model was fitted with the out-of-sample (from the CV) predicted age as the dependent variable and chronological age as the independent variable using the training data. The predicted age in the test set was corrected by subtracting the resulting intercept and dividing by the slope.

### 2.6 Correlation with cognitive measures

To understand the effect of bias correction and the impact of covariates on delta-behavior correlations, we performed correlations of behavior/cognitive measures from CamCAN and eNKI datasets (see section 2.1.2) with (1) uncorrected delta, (2) uncorrected delta with age as a covariate, (3) corrected delta, and (4) corrected delta with age as a covariate. If the bias correction eliminates the antagonistic relation between delta and age, we expect 2, 3, and 4 to give similar correlations. Furthermore, to assess the impact of data used for training, we performed these analyses using delta obtained from within-site and cross-site predictions.

### 2.7 Brain-age in clinical samples

Next, we used the ADNI dataset (Jack *et al*., 2008; Petersen *et al*., 2010) to validate our best-performing workflow on clinical samples. We estimated and compared the delta between CN, EMCI, LMCI, and AD subjects (Table 1d).

Our best-performing workflow trained on the four datasets was used to obtain the predictions, followed by application of bias correction model (see section 2.5). We compared two bias correction models, one derived using the CV predictions from the four training datasets and another using CN samples in ADNI data (Franke and Gaser, 2012). The group-wise corrected delta was compared using analysis of variance (ANOVA) followed by Bonferroni correction to counteract multiple comparisons. Emulating the scenario that application sites might have different numbers of CN samples, we learned bias correction models using CN sub-samples (0.1 to 0.9 fraction in steps of 0.1) drawn without replacement and applied them on the full CN and AD subjects. This process was repeated 100 times to estimate the variance of mean corrected delta in AD subjects.

Finally, we investigated associations between the corrected delta and three clinical test scores, MMSE, CDR, and FAQ. The correlations were computed using the whole sample and different diagnostic groups separately using Pearson’s correlation with age as a covariate for both sessions separately.

### 2.8 Comparison with brainageR

We compared the performance of our best-performing workflow with an already available brain-age estimation model, brainageR.

The brainageR model was trained on 3377 healthy individuals (age range: 18-92 years, mean ± SD age: 40.6 ± 21.4 years) from seven publicly available datasets using the GPR algorithm. It uses SPM12 to segment and normalize T1w images, from which GM, WM, and CSF vectors were extracted (using 0.3 probability masked brainageR-specific templates). PCA was used to reduce data dimensionality, and 435 components explaining 80% of the variance were retained. Note that brainageR uses three tissue types, while our focus is on GM.

We used (1) the OASIS-3 (N = 806; first scan per subject, age range: 43 to 89 years, mean ± SD age: 69.07 ± 9.06 years), and (2) the MyConnectome study (one subject scanned 20 times in a period of 3 years; age range: 45-48 years) datasets for this comparison. Additionally, we used sub-samples from OASIS-3 with test-retest durations of (1) less than 3 months (N = 36, age range = 43-81) and (2) between 3-4 years (N = 127, age range = 46-86) to evaluate test-retest reliability and longitudinal consistency, respectively (see section 2.4.3).

Next, to achieve a more direct comparison, we trained models with our best-performing workflow with brainageR’s preprocessing and tissue types. Following our focus on GMV, we compared; (1) CAT-preprocessed GMV (as done in the current study), (2) SPM-preprocessed GMV following brainageR, and (3) SPM-preprocessed GM, WM, and CSF images using brainageR. The latter investigates whether WM and CSF features provide complementary information leading to better predictions. For this, we evaluated within-site performance on IXI and CamCAN datasets (see section 2.4.1).

## 3. Results

### 3.1 Within-site predictions

The CV performance was obtained by averaging over 125 estimates–inner 5×5-fold CV, which was repeated 5 times in the outer CV (see section 2.4.1). The test predictions, i.e., single prediction per subject from the outer CV, were used to calculate the test performance. Finally, the CV and test performance were averaged separately over four datasets.

The average CV MAE (4.90-8.48 years) and the average test MAE (4.73-8.38 years) (Figure 2a, Table S2) were similar, indicating that the nested CV generalization estimates are indeed indicative of their test performance. The correlation between the true and predicted age on the test data ranged from 0.81-0.93, while the age bias (correlation between true age and delta) ranged from −0.22 to −0.83 (Table S2). Overall, all workflows showed a high similarity in their predictions (correlations 0.82-0.99 averaged across the four datasets; Figure S2). The top 20 workflows showed comparable CV and test MAE with a difference of less than 0.4 years.

**Figure 2.**
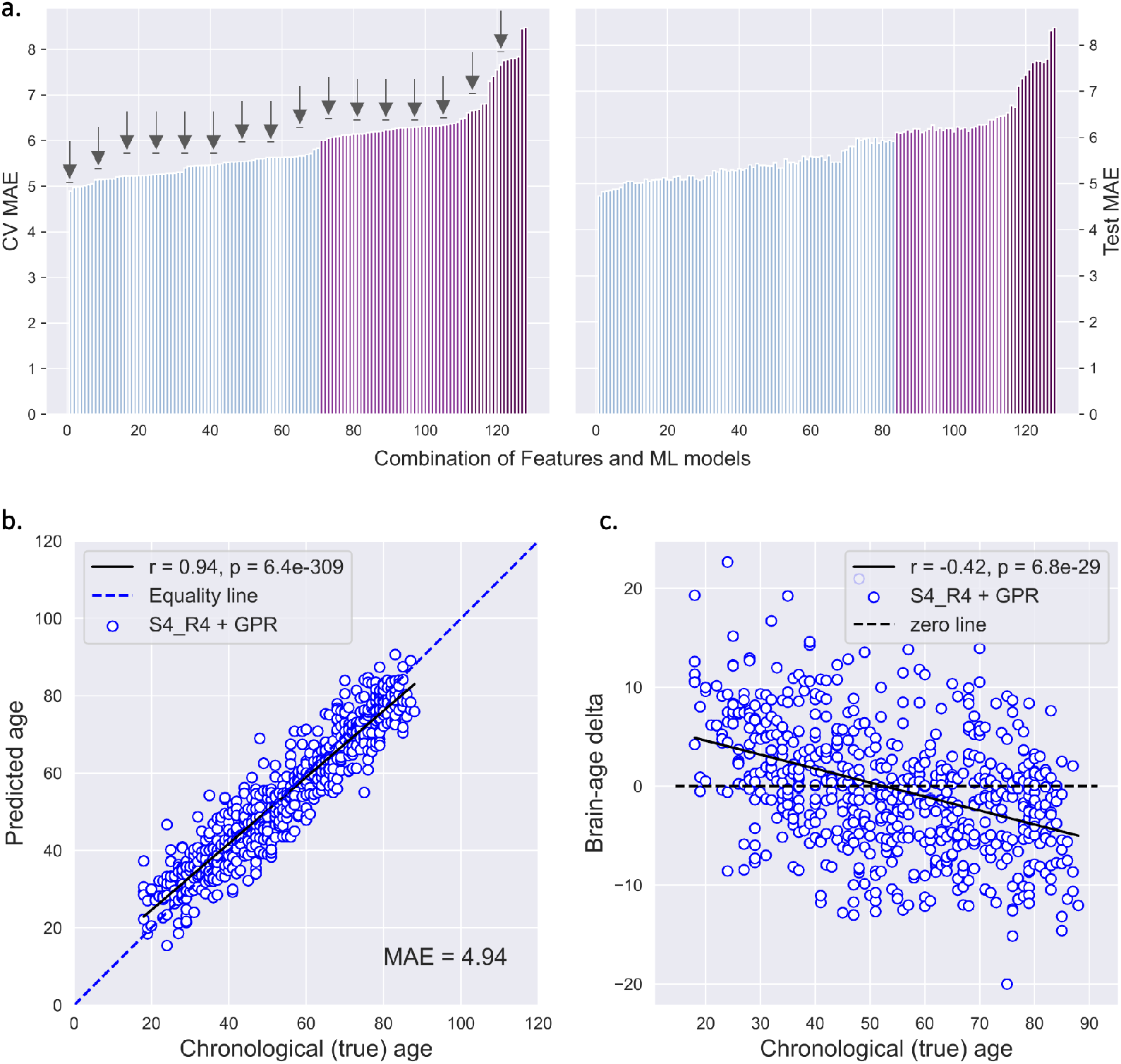
Within-site results. a. *(left)* CV MAE (averaged across four datasets) for 128 workflows arranged in increasing order (names of the workflow are given in Table S2) and *(right)* their corresponding averaged test MAE. Different colors of the bars indicate different MAE ranges (MAE < 6: blue, 6 >= MAE < 6.5: purple, MAE >= 6.5: dark purple). The arrows show the selected workflows for cross-site analysis. b. The scatter plot between the chronological age and within-site predicted age for the CamCAN data. The MAE and the correlation between predicted age and chronological age were MAE = 4.94 years and r = 0.94, p = 6.4e-309. c. The scatter plot between the chronological age and brain-age delta (predicted - chronological age) for CamCAN data. There was a negative correlation of r = −0.42, p = 6.8e-29 between brain-age delta and chronological age. The best-performing workflow from within-site predictions was S4_R4 + GPR.

Well-performing workflows primarily consisted of voxel-wise smoothed and resampled feature spaces with and without PCA, with S4_R4 (smoothed with a 4 mm FWHM kernel and resampled to 4 mm spatial resolution) generally performing better. Workflows with PCA often performed similarly to their respective non-PCA version. GPR, KRR, RR, and both RVR algorithms generally ranked high. Most algorithms performed worse with parcel-wise features, while RFR generally exhibited the worst performance.

The workflow S4_R4 + GPR performed the best (see Table 2a for its performance on each of the four datasets). This workflow showed the lowest average CV MAE with a high correlation between true and predicted age (Figure 2b) but a relatively high age bias (Figure 2c). The second-best workflow, S4_R4 + PCA + GPR, performed similarly to the best workflow. Other workflows with the S4_R4 feature space, with or without PCA, together with the KRR, RVRpoly, and RVRlin algorithms, performed comparably. From the 128 workflows, with a data-driven procedure, we selected 32 workflows while preserving diversity in workflows.

**Table 2.** The performance metric for the best workflow for different datasets. a. Within-site prediction (using S4_R4 + GPR) b. Cross-site prediction (using S4_R4 + PCA + GPR). Abbreviations: MAE: mean absolute error between true and predicted age, MSE: mean squared error between true and predicted age, R^2^: the proportion of variance of predicted age explained by the independent variables in the model, Corr (true, pred): Pearson’s correlation between true and predicted age, Age bias: Pearson’s correlation between true age and brain-age delta

| Datasets | N | a. Within-site results |  |  |  |  | b. Cross-site results |  |  |  |  |
| --- | --- | --- | --- | --- | --- | --- | --- | --- | --- | --- | --- |
| | | MAE | MSE | $R^2$ | Corr (true, pred) | Age bias | MAE | MSE | $R^2$ | Corr (true, pred) | Age bias |
| CamCAN | 651 | 4.94 | 39.54 | 0.89 | $r = 0.94$,<br>$p = 6.4e-309$ | $r = -0.42$,<br>$p = 6.8e-29$ | 4.75 | 38.35 | 0.89 | $r = 0.95$,<br>$p = 0.0e+00$ | $r = -0.23$,<br>$p = 3.1e-09$ |
| IXI | 562 | 4.76 | 35.20 | 0.87 | $r = 0.93$,<br>$p = 2.9e-252$ | $r = -0.48$,<br>$p = 3.5e-33$ | 6.08 | 57.35 | 0.79 | $r = 0.94$,<br>$p = 1.2e-267$ | $r = -0.18$,<br>$p = 2.2e-05$ |
| eNKI | 597 | 5.20 | 44.85 | 0.87 | $r = 0.93$,<br>$p = 8.1e-267$ | $r = -0.47$,<br>$p = 1.4e-33$ | 4.97 | 39.65 | 0.88 | $r = 0.94$,<br>$p = 9.7e-288$ | $r = -0.49$,<br>$p = 3.6e-38$ |
| 1000-BRAINS | 1143 | 4.04 | 26.65 | 0.83 | $r = 0.91$,<br>$p = 0.0e+00$ | $r = -0.50$,<br>$p = 2.0e-73$ | 5.13 | 41.03 | 0.73 | $r = 0.90$,<br>$p = 0.0e+00$ | $r = -0.15$,<br>$p = 2.0e-07$ |

### 3.2 Cross-site predictions

The 32 workflows selected from the within-site analysis included 23 workflows with voxel-wise feature spaces (S4_R4, S4_R8, S0_R4, S8_R4, S8_R8, S0_R8) with and without PCA and different ML algorithms (Figure 3a, Table S3). The remaining nine workflows included parcel-wise features (173, 473, 873, 1273) with GPR, RVRpoly, LR, ENR, and RFR algorithms. As expected, the average CV (5×5-fold on training data) MAE (4.28-7.39 years) was lower than the test (hold-out site) MAE (5.23-8.98 years). The test-set correlation between true and predicted age ranged from 0.82 to 0.93, while the age bias ranged from −0.27 to −0.75 (Table S3). The workflows that performed well within-site also performed well in cross-site predictions (Figure S6). These results indicate that the corresponding models could generalize well to data from a new unseen site.

**Figure 3.**
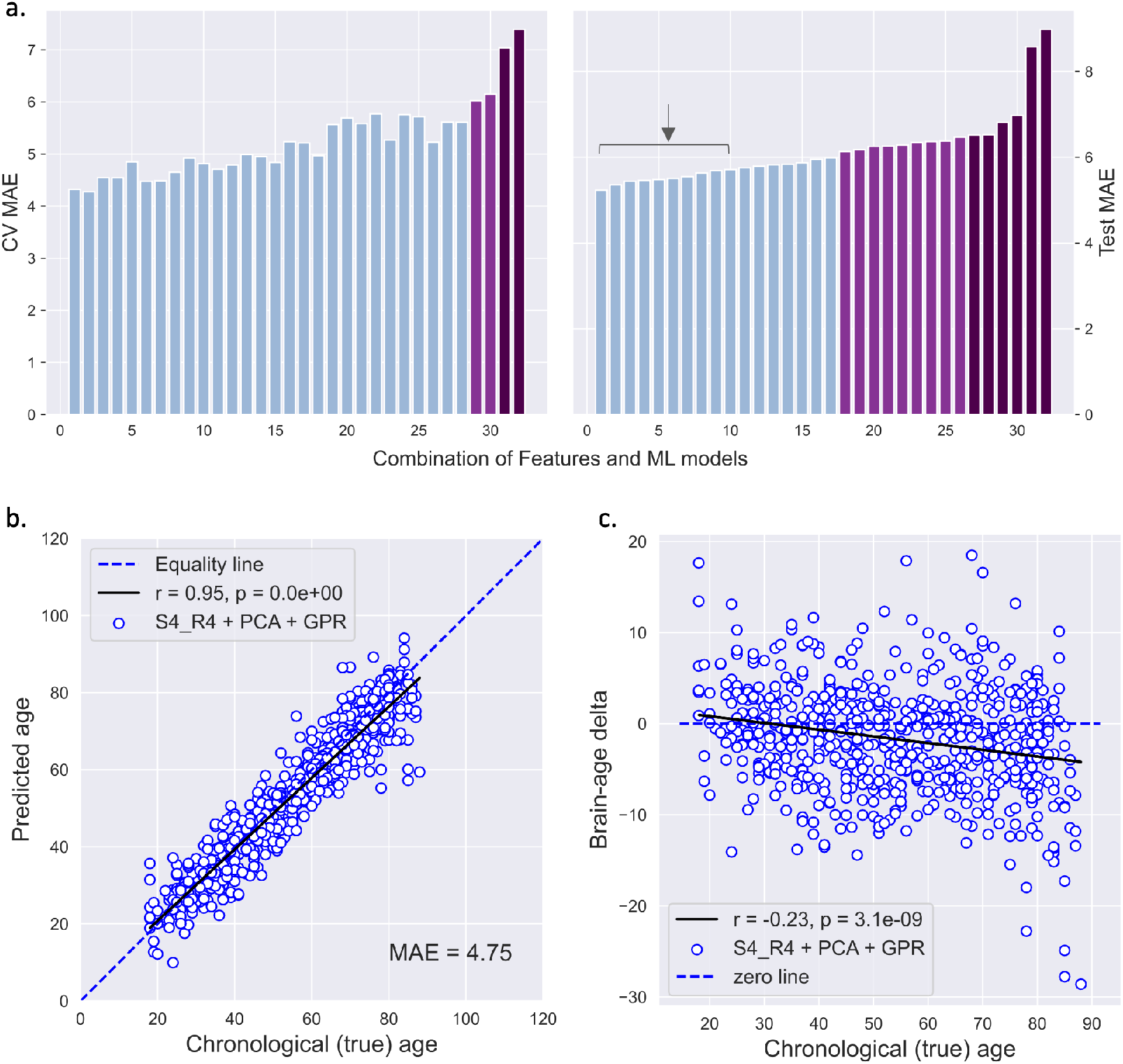
Cross-site results. a. *(left)* CV MAE (averaged across four runs) and *(right)* their corresponding averaged test MAE for the 32 selected workflows arranged in increasing order of the test MAE (names of the workflow are given in Table S3). Different colors of the bars indicate different MAE ranges (MAE < 6: blue, 6 >= MAE < 6.5: purple, MAE >= 6.5: dark purple). The arrow with the line shows the 10 selected workflows for test-retest reliability and longitudinal consistency analysis. b. The scatter plot between the chronological age and cross-site predicted age for the CamCAN data. The MAE and the correlation between predicted age and chronological age were MAE = 4.75 years and r = 0.95, p = 0.0e+00. c. The scatter plot between the chronological age and brain-age delta (predicted - chronological age) for CamCAN data. There was a negative correlation of r = −0.23, p = 3.1e-09 between brain-age delta and chronological age. The best-performing workflow from cross-site predictions was S4_R4 + PCA + GPR.

The top 10 workflows with low average test MAE consisted of only voxel-wise feature spaces (S4_R4, S4_R8, and S0_R4) with and without PCA. The ML algorithms included GPR, RVRlin, RR, and LR. The best-performing workflow was the S4_R4 + PCA + GPR with the lowest average test MAE, a high correlation between true and predicted age (Figure 3b), and moderate age bias (Figure 3c, see Table 2b for its performance on all four datasets), followed by the S4_R4 + GPR workflow. We selected these top 10 workflows with the lowest test MAE for further analysis.

### 3.3 Test-retest reliability and longitudinal consistency

The test-retest reliability and longitudinal consistency of the top 10 workflows selected from the cross-site evaluation were evaluated using the CoRR and OASIS-3 datasets.

For the retest duration of less than three months, all 10 workflows showed high test-retest reliability (CoRR: CCC = 0.95-0.98, age range 20-84 years; OASIS-3: CCC = 0.77-0.86, 43-81 years). In the CoRR dataset, for the retest duration of 1-2 years, CCC ranged between 0.94-0.97 (age range 18-88 years) (Table 3). These results show that the age was reliably estimated by the selected workflows.

**Table 3.** Concordance correlation coefficient (CCC) between brain-age delta from two sessions at different test-retest durations and their respective mean absolute error (MAE) between true and predicted age for CoRR and OASIS-3 datasets for the top 10 workflows.

|  | CoRR dataset |  |  |  |  |  | OASIS-3 dataset |  |  |
| --- | --- | --- | --- | --- | --- | --- | --- | --- | --- |
| Retest duration | < 3 months<br>(N = 86) |  |  | 1 – 2 years<br>(N = 95) |  |  | < 3 months<br>(N = 36) |  |  |
| Age range (years) | 20.0 - 84.0 |  |  | 18.0 - 88.0 |  |  | 42.66 - 80.90 |  |  |
| Workflows | MAE<br>(ses-1) | MAE<br>(ses-2) | CCC | MAE<br>(ses-1) | MAE<br>(ses-2) | CCC | MAE<br>(ses-1) | MAE<br>(ses-2) | CCC |
| S4_R4 + PCA + GPR | 4.808 | 5.008 | 0.97 | 4.374 | 4.204 | 0.95 | 4.2 | 3.801 | 0.80 |
| S4_R4 + GPR | 4.928 | 5.112 | 0.97 | 4.738 | 4.49 | 0.96 | 4.24 | 3.935 | 0.82 |
| S4_R4 + PCA + RVRlin | 5.811 | 5.757 | 0.97 | 5.156 | 5.072 | 0.96 | 5.288 | 5.223 | 0.83 |
| S4_R4 + RVRlin | 5.815 | 5.76 | 0.97 | 5.141 | 5.065 | 0.96 | 5.234 | 5.177 | 0.83 |
| S4_R8 + RVRlin | 6.375 | 6.265 | 0.95 | 5.444 | 5.33 | 0.96 | 5.109 | 5.2 | 0.77 |
| S4_R4 + RR | 5.64 | 5.653 | 0.98 | 5.174 | 5.277 | 0.97 | 4.918 | 4.71 | 0.85 |
| S4_R4 + PCA + RR | 5.742 | 5.732 | 0.98 | 5.288 | 5.404 | 0.97 | 4.988 | 4.744 | 0.85 |
| S0_R4 + LR | 6.281 | 6.359 | 0.96 | 6.251 | 6.293 | 0.94 | 4.949 | 5.161 | 0.86 |
| S4_R8 + LR | 6.763 | 6.676 | 0.97 | 6.497 | 6.434 | 0.97 | 5.811 | 5.896 | 0.79 |
| S4_R8 + RR | 6.232 | 6.185 | 0.97 | 5.975 | 6.016 | 0.97 | 5.332 | 5.328 | 0.81 |

Next, we evaluated the longitudinal consistency as the correlation between the difference in the predicted age and the difference in the chronological age using two scans at longer retest duration (Figure 4, Table S4). Six workflows out of 10 showed a significant positive linear relationship at the retest duration of 2-3.25 years (r between 0.451-0.437, p < 0.05) in the CoRR dataset. These workflows included the S4_R4 feature space with and without PCA with the GPR, RVRlin, and RR algorithms. In contrast, none of the workflows showed a linear relationship in the OASIS-3 dataset (retest duration 3-4 years).

**Figure 4.**
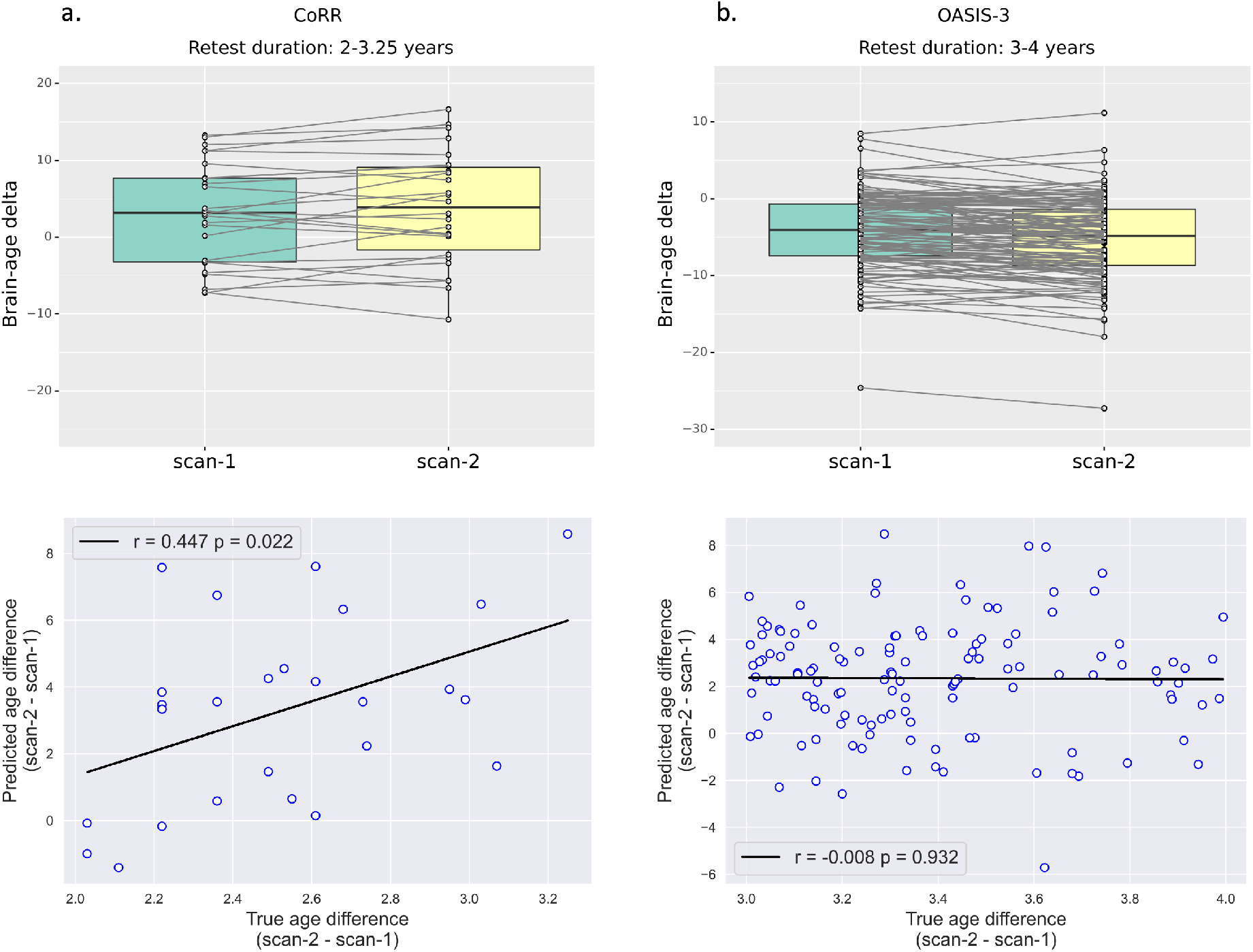
Longitudinal consistency. *(top)* The brain-age delta from two scans of the same subjects and *(bottom)* the scatter plot between the difference in chronological age and the difference in predicted age between two scans acquired within a retest duration of a. 2-3.25 years (CoRR dataset) b. 3-4 years (OASIS-3 dataset).

Although the workflows showed similar test-retest reliability and longitudinal consistency, the workflow S4_R4 + PCA + GPR showed the lowest MAE on these sub-samples (Table 3, Table S4). Therefore, considering all the analysis scenarios, within-site, cross-site, test-retest reliability, and longitudinal consistency, although other workflows were also competitive, we deemed the S4_R4 + PCA + GPR workflow as well-performing and chose it for further analysis.

### 3.4 Bias correction and correlation with behavioral/cognitive measures

In the CamCAN data, FI was negatively correlated with age (r = −0.661, p = 1.92e-80), while motor learning reaction time was positively correlated with age (r = 0.544, p = 1.11e-24). In the eNKI data, CWIT inhibition trial completion time (r = 0.361, p = 6.50e-12) and TMT number-letter switching trial completion time (r = 0.279, p = 1.45e-07) were positively correlated with age. On the other hand, WASI matrix reasoning scores were negatively correlated (r = −0.240, p = 6.03e-06), and WASI similarities scores were not correlated (r = 0.052, p = 0.332) with age (Table 4).

**Table 4.** Correlation of brain-age delta with various behavioral measures with and without bias correction. a. From within-site predictions. b. From cross-site predictions. Age was used as a covariate. Abbreviations: CWIT: Color-Word Interference Test, TMT: Trail Making Test, WASI-II: Wechsler Abbreviated Scale of Intelligence

| Dataset | Behavioral measure | N | Correlation with age | No bias correction |  | After bias correction |  |
| --- | --- | --- | --- | --- | --- | --- | --- |
|  |  |  |  | (a) No covariate | (b) With covariate | (c) No covariate | (d) With covariate |
| CamCAN | Fluid Intelligence (Cattel test) | 631 | $r = -0.661$,<br>$p = 1.9\text{e-}80$ | $r = -0.043$,<br>$p = 0.282$ | **$r = -0.154$,**<br>**$p = 0.0001$** | **$r = -0.157$,**<br>**$p = 7.2\text{e-}05$** | **$r = -0.154$,**<br>**$p = 0.0001$** |
| | Motor Learning (Reaction time) | 302 | $r = 0.544$,<br>$p = 1.1\text{e-}24$ | $r = 0.089$,<br>$p = 0.122$ | **$r = 0.181$,**<br>**$p = 0.002$** | **$r = 0.186$,**<br>**$p = 0.001$** | **$r = 0.180$,**<br>**$p = 0.002$** |
| eNKI | CWIT (Inhibition trial completion time) | 340 | $r = 0.361$,<br>$p = 6.5\text{e-}12$ | $r = 0.022$,<br>$p = 0.683$ | **$r = 0.109$,**<br>**$p = 0.045$** | $r = 0.094$,<br>$p = 0.084$ | **$r = 0.110$,**<br>**$p = 0.043$** |
| | TMT (Number-letter switching trial completion time) | 344 | $r = 0.279$,<br>$p = 1.5\text{e-}07$ | $r = -0.031$,<br>$p = 0.564$ | $r = 0.033$,<br>$p = 0.542$ | $r = 0.022$,<br>$p = 0.690$ | $r = 0.033$,<br>$p = 0.545$ |
| | WASI-II matrix reasoning | 347 | $r = -0.240$,<br>$p = 6.0\text{e-}06$ | $r = 0.026$,<br>$p = 0.627$ | $r = -0.030$,<br>$p = 0.581$ | $r = -0.019$,<br>$p = 0.728$ | $r = -0.029$,<br>$p = 0.590$ |
| | WASI-II similarities | 347 | $r = 0.052$,<br>$p = 0.332$ | $r = -0.033$,<br>$p = 0.536$ | $r = -0.022$,<br>$p = 0.685$ | $r = -0.023$,<br>$p = 0.667$ | $r = -0.021$,<br>$p = 0.698$ |

b. From cross-site predictions
| Dataset | Behavioral measure | N | Correlation with age | No bias correction |  | After bias correction |  |
| --- | --- | --- | --- | --- | --- | --- | --- |
|  |  |  |  | (a) No covariate | (b) With covariate | (c) No covariate | (d) With covariate |
| CamCAN | Fluid Intelligence (Cattel test) | 631 | $r = -0.661$,<br>$p = 1.9\text{e-}80$ | $r = 0.071$,<br>$p = 0.074$ | $r = -0.073$,<br>$p = 0.066$ | $r = -0.053$,<br>$p = 0.180$ | $r = -0.073$,<br>$p = 0.066$ |
| | Motor Learning (Reaction time) | 302 | $r = 0.544$,<br>$p = 1.1\text{e-}24$ | $r = -0.023$,<br>$p = 0.689$ | $r = 0.092$,<br>$p = 0.110$ | $r = 0.083$,<br>$p = 0.151$ | $r = 0.092$,<br>$p = 0.110$ |
| eNKI | CWIT (Inhibition trial completion time) | 340 | $r = 0.361$,<br>$p = 6.5\text{e-}12$ | $r = 0.005$,<br>$p = 0.931$ | **$r = 0.208$,**<br>**$p = 0.0001$** | $r = 0.065$,<br>$p = 0.230$ | **$r = 0.208$,**<br>**$p = 0.0001$** |
| | TMT (Number-letter switching trial completion time) | 344 | $r = 0.279$,<br>$p = 1.5\text{e-}07$ | $r = -0.007$,<br>$p = 0.898$ | **$r = 0.147$,**<br>**$p = 0.006$** | $r = 0.039$,<br>$p = 0.469$ | **$r = 0.147$,**<br>**$p = 0.006$** |
| | WASI-II matrix reasoning | 347 | $r = -0.240$,<br>$p = 6.0\text{e-}06$ | $r = 0.077$,<br>$p = 0.150$ | $r = -0.045$,<br>$p = 0.400$ | $r = 0.043$,<br>$p = 0.425$ | $r = -0.045$,<br>$p = 0.400$ |
| | WASI-II similarities | 347 | $r = 0.052$,<br>$p = 0.332$ | $r = -0.098$,<br>$p = 0.068$ | $r = -0.083$,<br>$p = 0.122$ | $r = -0.096$,<br>$p = 0.073$ | $r = -0.083$,<br>$p = 0.122$ |

As several ways have been proposed to obtain the correlation between delta and behavior, e.g., using bias-corrected delta or using age as a covariate, we evaluated several alternatives (see section 2.6).

#### 3.4.1 Within-site predictions

Within-site hold-out test predictions, i.e., single prediction per subject, were derived using the chosen workflow (S4_R4 + PCA + GPR). The bias correction model was estimated using the CV predictions on the same dataset. In both datasets, there was no residual age bias after bias correction: CamCAN, r = −0.17, p = 1.13e-05 and r = 0.00, p = 0.999; and eNKI, r = −0.20 p = 4.53e-07 and r = 0.001, p = 0.986, before and after correction, respectively (Figure S3).

We first calculated the correlation between the uncorrected delta and behavioral measures using age as a covariate (Table 4a,b). In the CamCAN data, a higher delta was associated with lower FI (r = −0.154, p = 0.0001) and higher motor learning reaction time (r = 0.181, p = 0.002). In the eNKI data, a higher delta was associated with lower response inhibition and selective attention, as indicated by a higher CWIT inhibition trial completion time (r = 0.109, p = 0.045). There were no correlations between delta and intelligence scores (WASI matrix reasoning and similarities). The results with age, age^2^, and gender as covariates showed a similar trend (Table S5a).

Next, we repeated this analysis with the corrected delta (Table 4a,c) and expected results similar to the uncorrected delta with age as a covariate. We indeed found similar correlations with FI (r = −0.157 p = 7.24e-05) and motor learning reaction time (r = 0.186 p = 0.001) in the CamCAN data, but no significant correlation with CWIT inhibition trial completion time (r = 0.094, p = 0.084) in the eNKI data. The correlations using corrected delta with covariate were highly similar to uncorrected delta with covariate (Table 4a, b & d).

#### 3.4.2 Cross-site predictions

Cross-site predictions were derived for the CamCAN and eNKI datasets using the S4_R4 + PCA + GPR workflow trained on the IXI + eNKI + 1000BRAINS (N = 2302) and IXI + CamCAN + 1000BRAINS (N = 2356) datasets, respectively.

In the CamCAN data, the bias correction model was successful with age bias before and after correction r = −0.23, p = 3.06e-09 and r = −0.04, p = 0.263, respectively. However, the correction was not successful in the eNKI data; the age bias was r = −0.49, p = 3.62e-38 and = −0.35, p = 8.39e-19 before and after correction, respectively (Figure S3). This result indicates that the bias correction might not always work well when applied to cross-site data.

Using age as a covariate on the uncorrected delta, we did not find a significant delta-behavior correlation in the CamCAN data. In the eNKI data, a higher delta was associated with lower response inhibition and selective attention, as indicated by a higher CWIT inhibition trial completion time (r = 0.208, p = 0.0001) and lower cognitive flexibility indicated by a higher TMT completion time (r = 0.147, p = 0.006) (Table 4b,b). There were no correlations between delta and intelligence scores (WASI matrix reasoning and similarities). The results with age, age^2^, and gender as covariates showed a similar trend (Table S5b).

Since there was a residual correlation between corrected delta and age, the correlations with behavior without age as a covariate can be unreliable. We, therefore, do not discuss correlations of the corrected delta without age as a covariate, but they are reported in Table 4b,c for completeness. Additionally, as expected, the correlations using corrected delta with age as a covariate were similar to uncorrected delta with covariate (Table 4b, b & d).

### 3.5 Predictions in the ADNI sample

At timepoint 1, the mean uncorrected delta was −5.97 years in CN, −4.39 in EMCI, −3.57 in LMCI, and −2.13 in AD (Figure 5a). In other words, the model underestimated age. The slope and intercept derived from the bias correction model using the training data (CV predictions) could not entirely correct for the under-estimation and age bias (Figure 5b). Bias correction using the whole ADNI CN sample removed the bias (average delta, CN:0, EMCI: 0.85, LMCI: 2.09, AD: 4.47 years) (Figure 5c). ANOVA revealed that the corrected delta differed significantly across the groups (F = 12.94, p = 3.10e-08), and post-hoc t-tests revealed significant differences between AD and CN (p = 1.16e-08), EMCI (p = 1.87e-05), LMCI (p = 0.043), and CN and LMCI (p = 0.022) after Bonferroni correction. At timepoint 2, the pattern was similar to timepoint 1 but with higher corrected delta values (EMCI: 1.15 years, LMCI: 2.88, AD: 6.59 years) (Figure 5e-f, Table 5). These results demonstrate that our model could capture the range of normal structural variation related to age in healthy subjects and deviance in both MCI and AD patients.

**Table 5.** Prediction performance on the ADNI data from two timepoints using the best-performing (S4_R4 + PCA + GPR) workflow. Abbreviations: CN: cognitively normal, EMCI and LMCI: early and late mild cognitive impairment, AD: Alzheimer’s disease

| Time-point | ADNI sample | N | MAE | MSE | Corr (true, pred) | Mean delta | Corrected mean delta (train samples) | Corrected mean delta (ADNI-CN samples) |
| --- | --- | --- | --- | --- | --- | --- | --- | --- |
| 1 | CN | 209 | 6.56 | 61.19 | $r = 0.76, p = 4.67e-40$ | -5.97 | -5.18 | 0.00 |
| | EMCI | 237 | 5.76 | 52.30 | $r = 0.72, p = 1.07e-38$ | -4.39 | -3.78 | 0.85 |
| | LMCI | 127 | 5.56 | 46.52 | $r = 0.75, p = 4.30e-24$ | -3.57 | -2.86 | 2.09 |
| | AD | 125 | 5.18 | 44.29 | $r = 0.66, p = 5.00e-17$ | -2.13 | -1.20 | 4.47 |
| 2 | CN | 153 | 6.56 | 62.73 | $r = 0.73, p = 5.46e-27$ | -6.05 | -5.27 | 0.00 |
| | EMCI | 197 | 5.57 | 50.82 | $r = 0.73, p = 1.23e-34$ | -4.32 | -3.66 | 1.15 |
| | LMCI | 104 | 5.68 | 47.75 | $r = 0.72, p = 6.54e-18$ | -3.25 | -2.44 | 2.88 |
| | AD | 61 | 5.31 | 44.12 | $r = 0.59, p = 6.09e-07$ | -0.76 | 0.31 | 6.59 |

**Figure 5.**
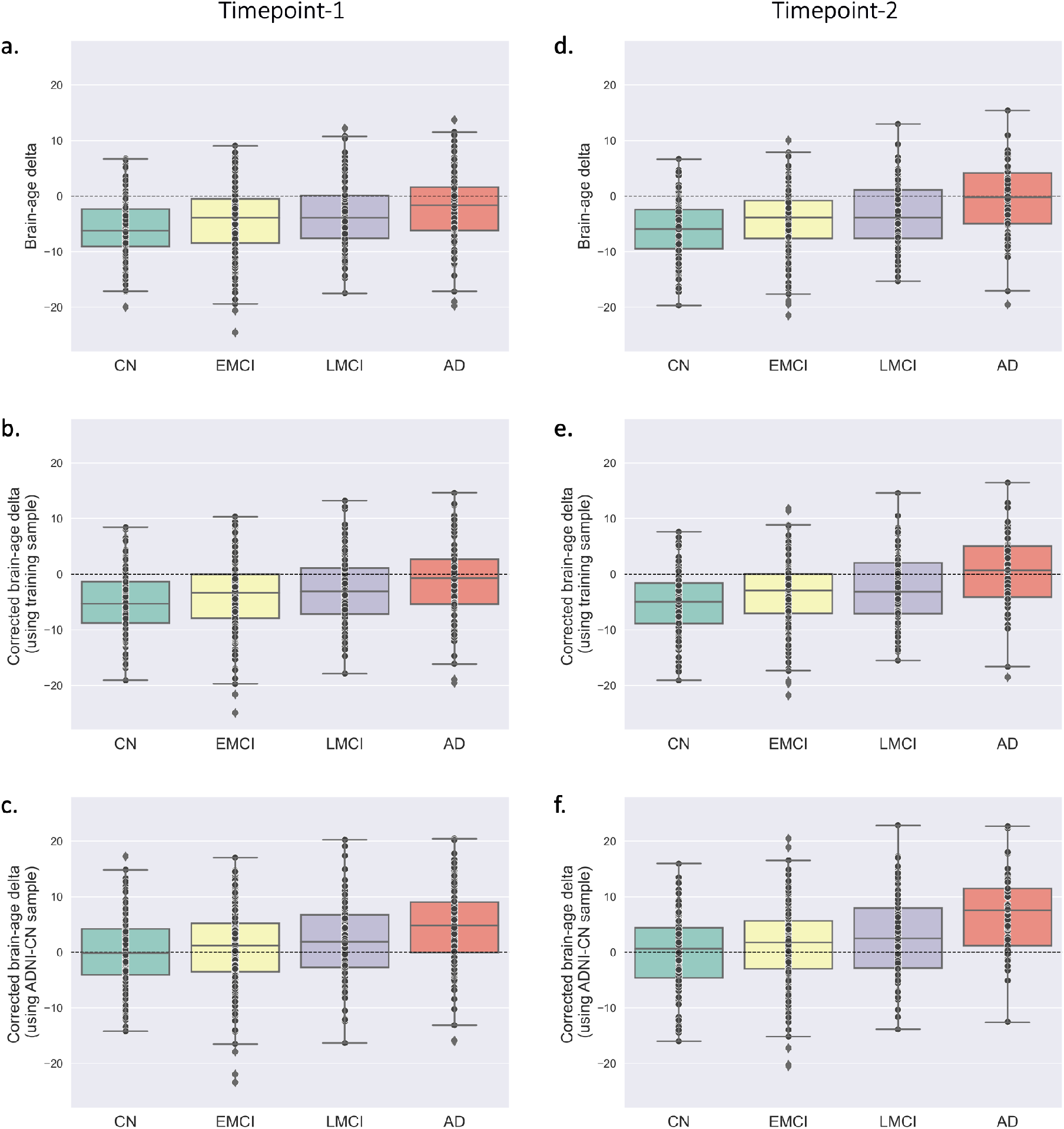
Brain-age delta in the clinical population. The box plot compares the delta between cognitive normal (CN), early mild cognitive impairment (EMCI), late mild cognitive impairment (LMCI), and Alzheimer’s disease (AD) from the ADNI sample at *(left)* timepoint-1 and *(right)* timepoint-2. Box plot with a & d. uncorrected delta. b & e. corrected delta using the CV predictions from the training set. c & f. corrected delta using the predictions from CN-ADNI subjects.

The correlations between CN sample-corrected delta and various clinical test scores were calculated with age as a covariate (Table 6). At timepoint 1, the delta was negatively correlated with MMSE (r = −0.255, p = 0.016) and positively correlated with FAQ (r = 0.275, p = 0.005) in the entire sample. No correlations were found in individual diagnostic groups or could not be calculated due to insufficient score data. At timepoint 2, the delta was negatively correlated with MMSE (r = −0.303, p = 2.40e-12) and positively correlated with CDR (r = 0.270, p = 7.35e-10) and FAQ (r = 0.331, p = 2.31e-14) in the whole sample. In the AD group, the delta was positively correlated with FAQ (r = 0.298, p = 0.021) but not with MMSE or CDR. In the LMCI group, the delta was positively correlated with FAQ (r = 0.309, p = 0.002), negatively correlated with MMSE (r = −0.227, p = 0.022), and not correlated with CDR. In the EMCI group, the delta positively correlated with CDR (r = 0.153, p = 0.034) but not MMSE and FAQ scores. No correlations were found in the CN group. The correlations with age, age^2^, and gender as covariates were similar (Table S6).

**Table 6.** Pearson’s correlation coefficients between corrected brain-age delta using S4_R4 + PCA + GPR workflow and cognitive measures (MMSE, CDR, and FAQ) using age as a covariate from the ADNI sample. The correlations were computed for the whole sample and each diagnostic group (CN, EMCI, LMCI and AD) separately from two timepoints. Abbreviations: MMSE: Mini-Mental State Examination, CDR: Global Clinical Dementia Rating Scale, FAQ: Functional Assessment Questionnaire; CN: cognitively normal, EMCI and LMCI: early and late mild cognitive impairment, AD: Alzheimer’s disease

|  | Timepoint-1 |  |  | Timepoint-2 |  |  |
| --- | --- | --- | --- | --- | --- | --- |
|  | MMSE | CDR | FAQ | MMSE | CDR | FAQ |
| CN | N = 68 | N = 67 | N = 74 | N = 153 | N = 147 | N = 149 |
| | $r = -0.202, p = 0.101$ | $r = 0.025, p = 0.841$ | $r = 0.153, p = 0.196$ | $r = -0.065, p = 0.427$ | $r = -0.019, p = 0.819$ | $r = 0.070, p = 0.399$ |
| EMCI | N = 3 | N = 3 | N = 3 | N = 196 | N = 194 | N = 193 |
| | n.a. | n.a. | n.a. | $r = -0.079, p = 0.272$ | **$r = 0.153, p = 0.034$** | $r = 0.091, p = 0.211$ |
| LMCI | N = 2 | N = 2 | N = 2 | N = 103 | N = 102 | N = 103 |
| | n.a. | n.a. | n.a. | **$r = -0.227, p = 0.022$** | $r = 0.115, p = 0.253$ | **$r = 0.309, p = 0.002$** |
| AD | N = 17 | N = 17 | N = 26 | N = 61 | N = 61 | N = 61 |
| | $r = -0.435, p = 0.092$ | $r = 0.221, p = 0.412$ | $r = 0.244, p = 0.240$ | $r = -0.186, p = 0.155$ | $r = 0.218, p = 0.094$ | **$r = 0.298, p = 0.021$** |
| Whole sample | N = 90 | N = 89 | N = 105 | N = 513 | N = 504 | N = 506 |
| | **$r = -0.255, p = 0.016$** | $r = 0.114, p = 0.290$ | **$r = 0.275, p = 0.005$** | **$r = -0.303, p = 2.40e-12$** | **$r = 0.270, p = 7.35e-10$** | **$r = 0.331, p = 2.31e-14$** |

We also found that the size of CN sample used for bias correction considerably impacts the mean corrected delta in AD subjects (Figure S7). Specifically, with fewer CN subjects, the variance of the corrected delta in AD was much higher in both sessions, e.g., at the timepoint 1 when using 21 CN samples, the mean AD delta ranged between ∼1-12 years and converged to 4.47 years as the sub-samples approached the complete sample.

### 3.6 Comparison with brainageR

Next, we compared the performance of our model trained with S4_R4 + PCA + GPR workflow and the brainageR model using the OASIS-3 and MyConnectome datasets (Figure 6).

**Figure 6.**
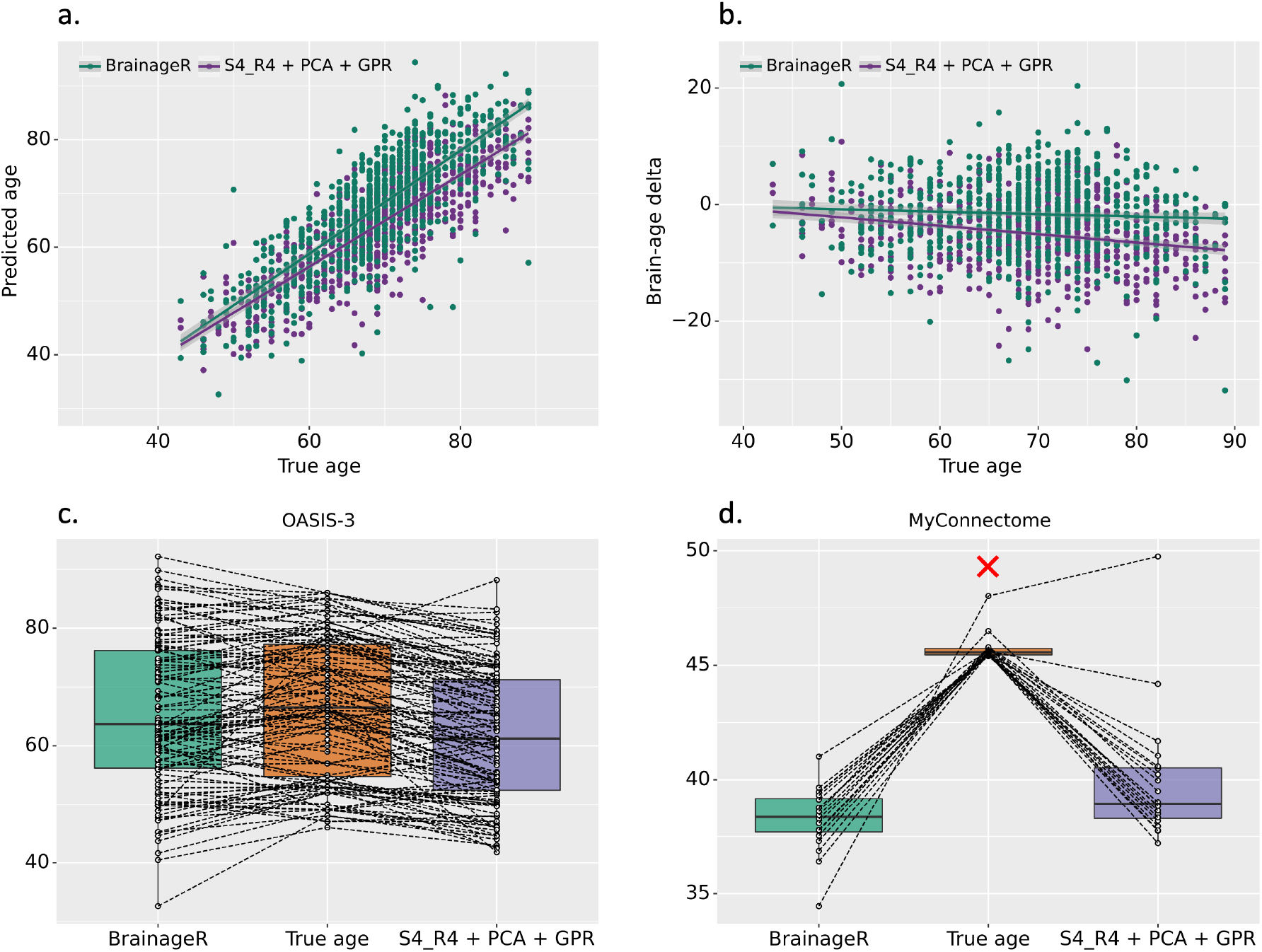
Comparison of our best workflow (S4_R4 + PCA + GPR) with the brainageR model. *(top)* The scatter plot between the chronological (true) age and a. the predicted age b. the brain-age delta from two models on OASIS-3 data. *(bottom)* The box plot comparing the predicted age from the two models on OASIS-3 data (for visual clarity, the plot is created using a random sub-sample; N = 120) d. MyConnectome data (the red cross indicates the outlier scan that was removed from the analysis; final N = 19).

brainageR (MAE = 5.07 years, age bias r = −0.058, p = 0.099) performed better than S4_R4 + PCA + GPR model (MAE = 5.96 years, age bias r = −0.238, p = 8.24e-12) on the OASIS-3 dataset (Figure 6a,b). The correlations between predicted ages and deltas from the two models were r = 0.844, p = 6.96e-220, and r = 0.535, p = 6.10e-61, respectively. There was a significant difference in the predicted age from the two models (paired t-test: t = −16.41, p = 1.90e-52). The correlation between true and predicted age for our model (r = 0.824, p = 2.59e-200) was significantly higher than that from brainageR (r = 0.805, p = 7.96e-185) (Steiger’s Z test (Steiger, 1980) z = 1.807, p = 0.035). The age bias was significantly higher for our model (z = −5.364, p = 0). Test-retest reliability on a sub-sample of the OASIS-3 dataset (retest duration < 3 months) was higher for brainageR (CCC = 0.94 vs. 0.80 for S4_R4 + PCA + GPR). However, none of the models showed longitudinal consistency at a retest duration of 3-4 years.

On the other hand, S4_R4 + PCA + GPR workflow (MAE = 6.21) performed significantly better than brainageR (MAE = 7.18) on the MyConnectome dataset (paired t-test: t = 2.95, p = 0.009; Figure 6d). Note that one outlier scan (true age = 48) was excluded from this analysis (final N = 19).

We then evaluated the within-site performance of our workflow using brainageR’s SPM preprocessing on IXI and CamCAN datasets. On both datasets, CAT-derived GM features performed better (IXI: MAE = 4.85 years, age bias r = −0.21, p = 7.39e-07; CamCAN: MAE = 5.01, age bias r = −0.17, p = 1.14e-05) than SPM-derived GM features (IXI: MAE = 6.25, age bias r = −0.40, p = 1.61e-22; CamCAN: MAE = 5.82, age bias r = −0.30, p = 2.66e-15) (Table 7). As brainageR uses three tissue types, we evaluated our workflow by incorporating three tissue types from the SPM preprocessing for a more direct comparison. This model performed better (IXI: MAE = 5.08, age bias r = −0.27, p = 1.64e-10; CamCAN: MAE = 4.88, age bias r = −0.25, p = 1.53e-10) than using only SPM-derived GM features, indicating that different tissue types carry complementary information (Table 7).

**Table 7.** Comparison of within-site performance between models trained with CAT-preprocessed GM features (*S*4_*R*4 + *PCA* + *GPR*; our framework), SPM-preprocessed GM features (*S*4_*R*4_*SPM*_ + *PCA* + *GPR*) and SPM-preprocessed GM+WM+CSF features 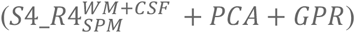 on IXI and CamCAN data. Abbreviations: MAE: mean absolute error, MSE: mean squared error, Corr (true, pred): Pearson’s correlation between true age and predicted age, Age bias: Pearson’s correlation between true age and brain-age delta

|  | Workflow | MAE | MSE | Corr (true, pred) | Age bias |
| --- | --- | --- | --- | --- | --- |
| **IXI**<br>(N = 562) | $S4\_R4 + PCA + GPR$ | 4.85 | 36.89 | $r = 0.93$, $p = 1.03e-247$ | $r = -0.21$, $p = 7.39e-07$ |
| | $S4\_R4_{SPM} + PCA + GPR$ | 6.25 | 62.34 | $r = 0.88$, $p = 1.15e-181$ | $r = -0.40$, $p = 1.61e-22$ |
| | $S4\_R4_{SPM}^{WM+CSF} + PCA + GPR$ | 5.08 | 40.80 | $r = 0.92$, $p = 3.98e-234$ | $r = -0.27$, $p = 1.64e-10$ |
| **CamCAN**<br>(N = 650) | $S4\_R4 + PCA + GPR$ | 5.01 | 40.89 | $r = 0.94$, $p = 6.45e-307$ | $r = -0.17$, $p = 1.14e-05$ |
| | $S4\_R4_{SPM} + PCA + GPR$ | 5.82 | 56.83 | $r = 0.92$, $p = 3.87e-258$ | $r = -0.30$, $p = 2.66e-15$ |
| | $S4\_R4_{SPM}^{WM+CSF} + PCA + GPR$ | 4.88 | 39.77 | $r = 0.94$, $p = 8.29e-308$ | $r = -0.25$, $p = 1.53e-10$ |

## 4. Discussion

### 4.1 Effect of feature space and ML algorithm

The wide range of options available for designing brain-age estimation workflows makes it challenging to disentangle the effect of feature space and ML algorithms. To this end, we investigated 128 workflows constituting combinations of 16 feature representations (voxel-wise and parcel-wise) extracted from GMV images and eight ML algorithms.

Previous studies have shown that the age prediction MAE ranges between ∼5-8 years for broad age range data (18-90 years) when using GMV features (Table S1). Our workflows showed performance in a similar range, with some of the workflows generalizing well to data from a new site. Specifically, the MAE ranged between 4.90-8.48 years in CV and 4.73-8.38 years in test data for within-site analysis and for cross-site analysis between 4.28-7.39 years and 5.23-8.98 years in CV and test data, respectively. The workflows showed high positive correlations between chronological age and predicted age for within-site (r between 0.81-0.93) and cross-site (r between 0.82-0.93) analyses. The workflows that performed well in within-site analysis also performed well in cross-site analysis. The lower cross-site CV MAE (4.28-7.39 years) compared to within-site CV MAE (4.90-8.48 years) might be because of the larger sample sizes in the cross-site analysis. This corroborates previous studies showing lower errors with larger training sets (Baecker *et al*., 2021*a*; de Lange *et al*., 2022). The age range of the training and test data affects the performance estimates. Specifically, when using a narrow age range, performance metrics such as MAE and RMSE are usually better than broad age range evaluations (Cole, 2020; Peng *et al*., 2021; de Lange *et al*., 2022). However, the lower errors and hence smaller brain-age delta values in those cases are not necessarily due to better model performance but rather because the predictions are closer to the mean age of the group. Here, our focus was on broad age range models, and the errors we obtained are within the range of what has been previously shown.

Our results showed that the choice of feature space and the ML algorithm both affect the prediction error. In general, feature spaces derived from voxel-wise GMV such as S4_R4, S4_R8, and S0_R4 in combination with GPR, KRR, RVRpoly, and RVRlin algorithms performed well in the within-site analysis. The results were similar when performing PCA while retaining 100% variance. Some of these selected workflows also performed well on cross-site analysis. Specifically, the voxel-wise GMV features smoothed with a 4 mm FWHM kernel and resampled to a spatial resolution of 4 mm, without and with PCA (S4_R4 and S4_R4 + PCA) together with the GPR algorithm performed best in both the within-site and cross-site analyses. A previous study has reported a voxel size of 3.73 mm^3^ and a smoothing kernel of 3.68 mm as the optimal parameters for processing GM images for brain-age prediction with a performance similar to our workflows (Lancaster *et al*., 2018). In general, parcel-wise features performed worse than voxel-wise features irrespective of the ML algorithm used, suggesting that the GMV summarized from parcels leads to a loss of age-related information. Our results align with a recent study comparing several ML models (GPR-dot product kernel, RVR-linear kernel, and SVR-linear kernel) trained on region-based and voxel-based features with or without PCA on a narrower age range (47–73 years) (Baecker *et al*., 2021*a*). They found minimal differences in performance due to the ML algorithms with voxel-based features performing better than region-based features.

Our results also indicate that the non-linear algorithm (GPR with RBF kernel) and the kernel-based algorithms (KRR and RVR) outperformed linear algorithms such as RR and LR. Surprisingly, the non-linear RFR algorithm performed the worst irrespective of the feature space used (Figure S4). This suggests that capturing distributional information using the RBF kernel, as we did using GPR, and use of kernels that capture the similarity between the GMV features in an invariant manner (e.g., Pearson correlation) is beneficial. These results corroborate a recent study that comprehensively evaluated 22 regression algorithms (test MAE between 4.63-7.14 years) in broad age range data (18-94 years) using GMV features and found SVR, KRR, and GPR with a diverse set of kernels to perform well (Beheshti *et al*., 2022).

In sum, the smoothed and resampled voxel-wise data (such as S4_R4, S4_R8) with either a non-linear or a kernel-based algorithm (GPR with RBF kernel, KRR with polynomial kernel degree (1 or 2), and RVR with linear and polynomial degree 1 kernels) are well suited for brain-age estimation. Sometimes, especially with a large number of features, PCA might help improve performance (Franke *et al*., 2010, Baecker *et al*., 2021*a*). However, we found the performance of many workflows with and without PCA to be similar, likely because resampling the GM images already reduces the number of features. Therefore, one could use the non-PCA version for immediate interpretability of the models; on the other hand, if computation is a constraint, then the PCA retaining 100% variance could be used without affecting the performance.

Future studies can investigate options to improve model generalizability, such as data harmonization to remove site effects and considerations for population structure (e.g., over-representative of the Caucasian population in the datasets used).

### 4.2 Test-retest reliability and longitudinal consistency

The brain-age estimates must be reliable within a subject. We found the delta to be reliable over a short scan delay (CoRR: CCC = 0.95-0.98, age range = 20-84; OASIS-3: CCC = 0.76-0.85, age range = 43-80). The reliability of delta within a short scan duration has been reported in previous studies. For example, one study showed an intraclass correlation coefficient (ICC) of 0.96 between deltas from subjects scanned an average of 28.35 ± 1.09 days apart (N = 20, mean age at first scan = 34.05 ± 8.71) (Cole *et al*., 2017). Another study showed an ICC of 0.93 in young adults from the OASIS-3 dataset (N = 20, age range = 19-34) scanned within a short delay of less than 90 days (Franke and Gaser, 2012). Another study found an ICC of 0.81 with a mean interval of 79 days between scans (N = 20, chronological age = 45 years) (Elliott *et al*., 2021).

Longitudinal consistency, i.e., chronologically proportionate increase in predicted age, is crucial for real-world application. We found a positive linear relationship between the difference in predicted age and the difference in chronological age at a retest duration of 2-3.25 years (N = 26; r = 0.447, p = 0.022) in the CoRR dataset. However, there was no correlation in the OASIS-3 dataset with a retest duration of 3-4 years (N = 127; r = −0.008, p = 0.932). Thus, the evidence of longitudinal consistency was weak. This can be speculatively explained by the maximum test-retest duration of 3-4 years which lies within the range of the MAE for the OASIS-3 dataset (MAE session-1: 5.08 and session-2: 5.86 years). Taken together, the high reliability supports the use of brain-age in clinical settings; however, further evaluations are needed to establish longitudinal consistency.

### 4.3 Effect of bias correction

Most brain-age estimation workflows produce biased results, i.e., over-estimation at younger ages and under-estimation at older ages (Liang *et al*., 2019). Therefore, correcting this age bias is important to facilitate individual-level decisions. Here, we adopted a bias correction model that does not use the chronological age of test samples for correction (Cole, 2020), as using chronological age can hamper fair comparison between workflows (de Lange *et al*., 2022).

The tested workflows generally showed negative associations between chronological age and delta for both within-site (r between −0.22 to −0.83) and cross-site (r between −0.27 to −0.75) predictions. However, this age bias was less pronounced in more accurate models (Figure S5). This result is in line with the previous work (de Lange *et al*., 2022) that showed that if input features are not informative enough to predict age, predictions will be closer to the median or mean age, leading to overestimation in younger subjects and underestimation in older subjects. Additionally, we found that the data used to estimate the bias correction models can significantly impact the corrected delta. Specifically, when using within-site data, the ensuing models corrected the age bias more adequately than when cross-site data were used (Figure S3). This discrepancy might be due to the difference in data properties, e.g., scanner-specific idiosyncrasy, between the training and the out-of-site test data. Our results suggest that a bias correction model might not always work well when applied to a new site, even when the training data itself consists of multiple sites. Consequently, using part of the test data to correct the age bias in the remaining test data works well (as seen in the ADNI data analysis, section 3.5). However, this might not be feasible when the test sample is small or in the extreme case, a single test subject is available.

How much data is needed for learning a bias correction model is an important but unexplored question. We investigated this by learning bias correction models from sub-samples of the CN subjects from ADNI data. Smaller samples led to higher variance in the efficacy of bias correction models when applied to AD patients (Varoquaux *et al*., 2017). For instance, at the smallest sample size (N = 21), the average corrected delta of the AD patients varied from 1 to 12 years (Figure S7, ADNI timepoint 1). It is likely that different studies use different samples for bias correction, so the results should be interpreted and compared with caution. This result shows the importance of using large samples for bias correction and emphasizes careful analysis and reporting of the results.

### 4.4 Correlation with behavior

The correlation of delta with behavioral measures is sensitive to whether the delta was adjusted for age, either via bias correction or using it as a covariate. For instance, the uncorrected delta was not correlated with FI and motor learning reaction time (in CamCAN data) or CWIT inhibition trial completion time (in eNKI data); however, significant correlations were obtained using age-adjusted delta. Thus, it is important to control for age when analyzing correlations between delta and behavioral measures.

Using out-of-sample predictions from within-site analysis, we found that a higher uncorrected delta (with age as a covariate) was associated with lower FI, higher motor learning reaction time (from CamCAN data), and lower response inhibition and selective attention, indicated by higher CWIT inhibition trial completion time (from eNKI data). We expected these correlations to be similar to correlations calculated using corrected delta (de Lange and Cole, 2020), as there was no significant age bias. In the CamCAN data, the behavioral correlations using uncorrected delta with age as a covariate and corrected delta were quite similar (FI: r = −0.154, p = 0.0001 vs. r = −0.157, p = 7.24e-05; motor learning reaction time: r = 0.181, p = 0.002 vs. r = 0.186, p = 0.001). However, the correlation of CWIT inhibition trial completion time with uncorrected delta with age as a covariate was significant but not when using the corrected delta (r = 0.109, p = 0.045 vs. r = 0.094, p = 0.084). This slight difference could potentially be explained by the small effect size and differences inherent in the two methods used for correction.

We also found that there was disagreement between delta-behavior correlations from within-site and cross-site predictions with age as a covariate. For instance, CamCAN showed significant correlations with FI and motor learning reaction time with within-site delta but not with cross-site delta. On the other hand, eNKI showed significant correlations only with CWIT inhibition trial completion time using within-site delta, but a significant correlation with TMT completion time was also found using cross-site delta. These results indicate that the subtle differences in predictions can impact behavioral correlations, even though the two predictions were highly correlated (CamCAN: r = 0.961, eNKI: r = 0.962; Figure S6). Thus, the delta-behavior correlations, whether using within-site or cross-site delta, should be interpreted with caution.

Taken together, within-site data yields better bias correction models, as we observed in two scenarios, behavioral correlations and delta estimation. However, when enough data are not available, the resulting models may fail to correct the age bias, leading to high variability in the mean delta (Figure S7). We therefore caution the practitioners and recommend carefully assessing bias correction models, e.g., using bootstrap analysis, before application. We observed that subtle differences in predicted age (within-site vs. cross-site) lead to different behavioral correlations, which can question the impact of the workflow used for prediction, the analysis method used for computing behavioral correlation (corrected delta versus covariates) and their interaction. Future studies should focus on disentangling such intricacies before applying the brain-age paradigm in practice.

### 4.5 Higher brain-age delta in neurodegenerative disorders

Neurodegenerative disorders such as AD, MCI, and Parkinson’s disease (PD) are accompanied by brain atrophy. Many studies have shown a decrease in global and local GMV in MCI and AD (Good *et al*., 2001; Karas *et al*., 2004; Fjell *et al*., 2014) and also in a broad range of neuropsychiatric disorders (Kaufmann *et al*., 2019). Consequently, an increased delta, i.e., older appearing brains, has been reported in patients with MCI (3-8 years) and AD (∼10 years) (Franke and Gaser, 2012; Gaser *et al*., 2013; Varikuti *et al*., 2018). We assessed the delta in CN, EMCI, LMCI, and AD patients by applying our best-performing workflow followed by a bias correction model estimated on CN. We found that brain aging is advanced by ∼4.5-7 years in AD, ∼2-3 years in LMCI, and ∼1 year in EMCI (timepoint 1-timepoint 2). Furthermore, the delta was correlated with measures associated with disease severity and cognitive impairment in MCI and AD patients. Thus, in line with previous studies, brain-age delta confirmed its potential to indicate accelerated brain aging in neurodegenerative diseases based on structural MRI data (Franke and Gaser, 2012; Varikuti *et al*., 2018; Cole *et al*., 2019, 2020; Eickhoff *et al*., 2021; Lee *et al*., 2021).

In the current study, we applied only one workflow to the ADNI data. However, it is important to note that different workflows can lead to different delta estimates (Beheshti *et al*., 2022) and, consequently, different correlations with cognitive measures. In addition, the mean corrected delta in the patient group depends on the type (within-site or cross-site) and size of sample used for bias correction. Thus, the results should be interpreted with caution when comparing different studies.

### 4.6 Comparison with brainageR and effect of preprocessing and tissue types

brainageR outperformed our workflow in the OASIS-3 sample (N = 806; MAE = 5.07 vs. MAE = 5.96) with lower age bias (r = −0.058, p = 0.099 vs. r = −0.238, p = 8.24e-12) and higher test-retest reliability in a sub-sample (N = 36; CCC = 0.94 vs. CCC = 0.80). However, it performed worse in the MyConnectome data (MAE = 7.18 vs. MAE = 6.21). The differences between brainageR and our best workflow might be driven by differences in training data, preprocessing, and the use of three tissue types by brainageR as opposed to us using only GM. To investigate this further, we performed two additional analyses.

Different VBM tools can provide different GMV estimates, influencing the estimated association with age (Tavares *et al*., 2019). The CAT-derived GMV features performed better than brainageR’s SPM preprocessing (both with S4_R4 + PCA for feature extraction together with the GPR algorithm for learning) in terms of MAE (e.g., IXI: MAE = 4.85 vs. MAE = 6.25), the correlation between true and predicted age (r = 0.93 vs. r = 0.88, p < 1e-6) and age bias (r = −0.21 vs. r = −0.40, p < 1e-6) (Table 7). We further found that the predictions when using three tissue types from SPM (GM, WM, and CSF) were better (MAE = 5.08, r = 0.92, p < 1e-6, age bias: r = −0.27, p < 1e-6). This is in line with a previous study that showed a slight performance improvement when using both GM and WM compared to only GM (Cole et al., 2017a). Features from different tissue types may carry complementary information regarding age, providing better predictions and lower age bias. Many previous studies have used GM and WM together as features (Franke and Gaser, 2012, Cole et al., 2017b, 2018, 2020), and others have used all three tissue types (Monté-Rubio et al., 2018; Xifra-Porxas et al., 2021; Hobday et al., 2022). CAT-derived GMV performed similarly to SPM-derived three tissue types with slightly lower age bias for the former (Table 7), showing the suitability of GM for this task following its clinical relevance in neurodegenerative disorders (Karas et al., 2004; Wu et al., 2021).

## 5. Conclusion

Numerous choices exist for designing a workflow for age prediction. The systematic evaluation of different workflows on the same data in different scenarios (within-site, cross-site, and test-retest reliability) revealed a substantial impact of feature representation and ML algorithm choices. Notably, voxel-wise GM features, especially smoothed with a 4 mm FWHM kernel and resampled to a spatial resolution of 4 mm (S4_R4), were better than parcel-wise features. Additionally, performing PCA did not affect the prediction performance, but it can help reduce computational resources. ML algorithms, including Gaussian process regression with the radial basis kernel, kernel ridge regression with polynomial kernel degree 1 or 2, and relevance vector machine with linear and polynomial degree 1 kernels, performed well. Overall, some workflows performed well on out-of-site data and showed high test-retest reliability but only moderate longitudinal reliability. Consistent with the literature, we found a higher delta in Alzheimer’s and mild cognitive impairment patients after correcting the delta with a large sample of controls. Our results provide evidence for the future application of delta as a biomarker but also caution regarding analysis setup and data used for behavioral correlations and bias correction. Findings from the current study can serve as guidelines for future brain-age prediction studies.

## Ethics statement

Ethical approval and informed consent were obtained locally for each study covering both participation and subsequent data sharing. The ethics proposals for the use and retrospective analyses of the datasets were approved by the Ethics Committee of the Medical Faculty at the Heinrich-Heine-University Düsseldorf.

## Data/code availability statement

The codes used for preprocessing, feature extraction and model training are available at https://github.com/juaml/brainage_estimation.

## Acknowledgments

This study is supported by Deutsche Forschungsgemeinschaft (DFG, PA 3634/1-1 and EI 816/21-1), the National Institute of Mental Health, the Helmholtz Portfolio Theme “Supercomputing and Modelling for the Human Brain” and the European Union’s Horizon 2020 Research and Innovation Program grant agreement 945539 (HBP SGA3).

Data for the MyConnectome project were obtained from the OpenNeuro database (ds000031).

The clinical data used in the preparation of this article were obtained from the Alzheimer’s Disease Neuroimaging Initiative (ADNI) database (adni.loni.usc.edu). The ADNI was launched in 2003 as a public-private partnership led by Principal Investigator Michael W. Weiner, MD. The primary goal of the ADNI has been to test whether serial magnetic resonance imaging (MRI), positron emission tomography (PET), other biological markers, and clinical and neuropsychological assessment can be combined to measure the progression of mild cognitive impairment (MCI) and early Alzheimer’s disease (AD). For up-to-date information, see www.adni-info.org. Data collection and sharing for this project was funded by the Alzheimer’s Disease Neuroimaging Initiative (ADNI) (National Institutes of Health Grant U01 AG024904) and DOD ADNI (Department of Defense award number W81XWH-12-2-0012). ADNI is funded by the National Institute on Aging, the National Institute of Biomedical Imaging and Bioengineering, and through generous contributions from the following: AbbVie, Alzheimer’s Association; Alzheimer’s Drug Discovery Foundation; Araclon Biotech; BioClinica, Inc.; Biogen; Bristol-Myers Squibb Company; CereSpir, Inc.; Cogstate; Eisai Inc.; Elan Pharmaceuticals, Inc.; Eli Lilly and Company; EuroImmun; F. Hoffmann-La Roche Ltd. and its affiliated company Genentech, Inc.; Fujirebio; GE Healthcare; IXICO Ltd.; Janssen Alzheimer Immunotherapy Research & Development, LLC.; Johnson & Johnson Pharmaceutical Research & Development LLC.; Lumosity; Lundbeck; Merck & Co., Inc.; Meso Scale Diagnostics, LLC.; NeuroRx Research; Neurotrack Technologies; Novartis Pharmaceuticals Corporation; Pfizer Inc.; Piramal Imaging; Servier; Takeda Pharmaceutical Company; and Transition Therapeutics. The Canadian Institutes of Health Research is providing funds to support ADNI clinical sites in Canada. Private sector contributions are facilitated by the Foundation for the National Institutes of Health (www.fnih.org). The grantee organization is the Northern California Institute for Research and Education, and the study is coordinated by the Alzheimer’s Therapeutic Research Institute at the University of Southern California. ADNI data are disseminated by the Laboratory for Neuro Imaging at the University of Southern California.

