## Supplementary Material for "Brain-age prediction: a systematic comparison of machine learning workflows"

**Table S1.** Comparison of various brain-age prediction studies using voxel-wise gray matter (GM) data with or without additional pre-processing. The test performance is reported on heathy population, * independent test set, # No cross-validation, - information not available

Abbreviations: CV: cross-validation, MAE: mean absolute error, RMSE: root mean squared error, HC: healthy controls, RVR: relevance vector regression, SVR: support vector regression, GPR: Gaussian process regression, LASSO: least absolute shrinkage and selection operator, ENR: elastic net regression, PCA: principal component analysis, ICA: independent component analysis, SCN: Structural covariance network

| | | | | **Cross-validation (CV)** | | | | | **Test set (Healthy)** | | | | |
|---|---|---|---|---|---|---|---|---|---|---|---|---|---|
| | **Author** | **Model** | **Dimensionality reduction** | **N** | **Age range** | **MAE** | **RMSE** | **Correlation** | **N** | **Age range** | **MAE** | **RMSE** | **Correlation** |
| 1 | (Ashburner, 2007) | RVR | - | 471 | 17-79 | - | 6.50 | 0.86 | - | - | - | - | - |
| 2 | (Franke *et al.*, 2010) | RVR | PCA | 410 | 19-86 | # | - | - | 137/ *108 | 19-86/ 20-59 | 4.61/ 5.44 | 5.90/ 6.73 | 0.94/ 0.89 |
| | | RVR | - | 410 | 19-86 | # | - | - | 137/ *108 | 19-86/ 20-59 | 4.96/ 5.57 | - | - |
| | | SVR | PCA | 410 | 19-86 | # | - | - | 137/ *108 | 19-86/ 20-59 | 4.85/ 5.42 | - | - |
| | | SVR | - | 410 | 19-86 | # | - | - | 137/ *108 | 19-86/ 20-59 | 4.85/ 5.51 | - | - |
| 3 | (Gaser *et al.*, 2013) | RVR | PCA | 320 | 51-94 | # | | | 64 | 51-83 | 3.8 | | |
| 4 | (Koutsouleris *et al.*, 2014) | SVR | PCA | 800 | 18-65 | 4.60 | - | | - | - | - | - | |
| 5 | (Su *et al.*, 2013) | RVR | sparse representation | 290 / 84 | 18-91 / 19-79 | 5.69 / 4.67 | - | | - | - | - | - | |
| 6 | (Cole *et al.*, 2015) | GPR | - | 1537 | 18-90 | 6.20 | - | 0.92 | *113 | 43.3 ± 20.24 | 5.80 | - | 0.93 |
| **7** | (Schnack *et al.*, 2016) | SVR | | 386 | 16-67 | 4.31 | - | - | *55 | 19-48 | 3.86 | - | - |

| | | | | | | | | | | | | | |
|---|---|---|---|---|---|---|---|---|---|---|---|---|---|
| 8 | (Cole *et al.*, 2017) | GPR | | 1601 | 18-90 | - | - | - | 200 | 18-90 | 4.66 | 6.01 | 0.95 |
| 9 | (Beheshti *et al.*, 2018) | SVR | feature- ranking strategy for feature selection | 385 | 63.50 ± 7.64 | - | - | - | *146 | 68.28 ± 5.61 | 4.02 | 5.10 | - |
| 10 | (Varikuti *et al.*, 2018) | LASSO | mixed | 1084 | 18-81 | 4.9 | - | 0.91 | - | - | - | - | - |
| | | | 1000b | 693 | 55-75 | 3.4 | - | 0.69 | - | - | - | - | - |
| | | | OPNMF - MIXED | 1084 | 18-81 | 6.1 | - | 0.88 | 693 (1000B) | * 55-75 | 6.1 | - | 0.55 |
| | | | OPNMF – 1000B | 693 | 55-75 | 3.6 | - | 0.65 | 239 (old mix) | * 55-75 | 3.7 | - | 0.53 |
| 11 | (Gutierrez Becker *et al.*, 2018) | GPR | PCA | 1543 | 6-92 | 5.65 | - | | - | - | - | - | |
| 12 | (Lancaster *et al.*, 2018) | SVR | resampling and smoothing | 1803 | 16-90 | - | - | | 200 | 16-90 | 5.08 | - | 0.94 |
| | | | resampling and smoothing | 2003 | 16-90 | - | - | | *648 | 18-88 | 6.08 | - | 0.93 |
| 13 | (Le *et al.*, 2018) | SVR | - | 475 | 18-60 | 5.1 | - | 0.82 | * 489 (incl. non-HC) | 18-56 | 4.84 | - | |
| 14 | (Monté-Rubio *et al.*, 2018) | GPR | | 562 | 20-86 | 5.1 | 6.4 | - | - | - | - | - | - |
| 15 | (Eavani *et al.*, 2018) | SVR | 583 ROIs | 400 | 50-96 | 4.41 | - | 0.80 | - | - | - | - | - |
| 16 | (Bagarinao *et al.*, 2018) | LASSO | ICA | 147 | 21-86 | - | - | - | 146 | 21-86 | 7.18 | | 0.86 |
| 17 | (Kuo *et al.*, 2020) | LASSO | ICA-SCN | 800 | 50-90 | 3.72 | - | 0.83 | 109 | 0-90 | 3.66 | - | 0.79 |

| | | | | | | | | | | | | | |
|---|---|---|---|---|---|---|---|---|---|---|---|---|---|
| 18 | (Boyle *et al.*, 2021) | Ensemble ENR | | 1359 | 18-88.36 | 7.28 | - | 0.85 | * 175 | 48-94 | 7.60 | - | 0.78 |
| | | | | | | | | | * 380 | 19-80 | 8.56 | - | 0.87 |
| | | | | | | | | | * 487 | 50-88 | 8.42 | - | 0.65 |
| 19 | (Baecker *et al.*, 2021) | SVR | - | 10480 | 47-73 | 4.33 | 5.43 | 0.73 | * 334 | 47-73 | 4.69 | 5.92 | 0.71 |
| | | RVR | - | 10480 | 47-73 | 3.69 | 4.60 | 0.75 | * 334 | 47-73 | 3.66 | 4.51 | - |
| | | SVR | PCA | 10480 | 47-73 | 3.89 | 4.86 | 0.71 | * 334 | 47-73 | 3.77 | 4.65 | 0.74 |
| | | RVR | PCA | 10480 | 47-73 | 3.90 | 4.85 | 0.71 | * 334 | 47-73 | 3.82 | 4.65 | 0.74 |
| | | GPR | PCA | 10480 | 47-73 | 3.90 | 4.85 | 0.71 | * 334 | 47-73 | 3.81 | 4.64 | 0.74 |
| 20 | (Eickhoff *et al.*, 2021) | Ensembled linear SVR | Atlas | 3960 | 30-95 | - | - | - | * 172 | 62.8 ± 8.3 | 4.4 | - | 0.84 |
| 21 | (Beheshti *et al.*, 2022) | 22 algorithms | resampling and PCA | 788 | 18-94 | 3.04-6.71 | 4.45-9.43 | - | 88 | 20-87 | 4.63-7.14 | 6.29-9.67 | - |

**Table S2.** Within-site results. Averaged CV MAE, test MAE and test correlation between predicted and true age for 128 workflows arranged in increasing order of CV MAE (averaged over four datasets). The 32 selected workflows for cross-site analysis are marked in bold letters with asterisk (*) .

| S.No. | Workflow Names | Average CV MAE | Average Test MAE | Averaged test correlation | Average test age-bias |
|---|---|---|---|---|---|
| **1** | **S4_R4 + GPR *** | **4.898** | **4.733** | **0.928** | **-0.468** |
| **2** | **S4_R4 + PCA + GPR *** | **4.978** | **4.822** | **0.928** | **-0.218** |
| 3 | S4_R4 + KRR | 4.99 | 4.838 | 0.928 | -0.407 |
| 4 | S4_R4 + PCA + RVRpoly | 4.992 | 4.85 | 0.928 | -0.386 |
| 5 | S4_R4 + RVRpoly | 5.008 | 4.871 | 0.928 | -0.398 |
| 6 | S4_R4 + PCA + KRR | 5.033 | 4.879 | 0.928 | -0.401 |
| 7 | S4_R8 + GPR | 5.056 | 4.916 | 0.926 | -0.451 |
| 8 | S4_R8 + PCA + GPR | 5.14 | 5.014 | 0.926 | -0.226 |
| **9** | **S4_R4 + RVRlin *** | **5.15** | **5.045** | **0.921** | **-0.426** |
| **10** | **S4_R4 + PCA + RVRlin *** | **5.15** | **5.047** | **0.921** | **-0.426** |
| 11 | S0_R4 + KRR | 5.158 | 5.005 | 0.926 | -0.486 |
| 12 | S0_R4 + PCA + RVRpoly | 5.161 | 5.009 | 0.926 | -0.486 |
| 13 | S0_R4 + RVRpoly | 5.163 | 5.015 | 0.926 | -0.486 |
| 14 | S4_R8 + KRR | 5.207 | 5.099 | 0.918 | -0.408 |
| 15 | S8_R4 + GPR | 5.216 | 5.05 | 0.921 | -0.433 |
| 16 | S0_R4 + PCA + GPR | 5.222 | 5.076 | 0.924 | -0.233 |
| **17** | **S4_R4 + RR *** | **5.223** | **5.084** | **0.918** | **-0.495** |
| **18** | **S4_R4 + PCA + RR *** | **5.224** | **5.09** | **0.918** | **-0.495** |
| 19 | S4_R8 + PCA + KRR | 5.226 | 5.123 | 0.918 | -0.389 |
| 20 | S0_R4 + PCA + KRR | 5.232 | 5.082 | 0.926 | -0.496 |
| 21 | S8_R8 + GPR | 5.238 | 5.074 | 0.921 | -0.431 |
| 22 | S4_R8 + RVRpoly | 5.24 | 5.145 | 0.916 | -0.393 |
| 23 | S4_R8 + PCA + RVRpoly | 5.245 | 5.159 | 0.916 | -0.369 |
| 24 | S4_R4 + ENR | 5.252 | 5.081 | 0.921 | -0.526 |
| **25** | **S8_R4 + PCA + GPR *** | **5.259** | **5.086** | **0.921** | **-0.291** |
| **26** | **S4_R8 + RVRlin *** | **5.268** | **5.168** | **0.916** | **-0.411** |
| 27 | S4_R8 + PCA + RVRlin | 5.268 | 5.168 | 0.916 | -0.411 |
| 28 | S8_R8 + PCA + GPR | 5.281 | 5.099 | 0.921 | -0.293 |
| 29 | S0_R4 + GPR | 5.282 | 5.071 | 0.924 | -0.586 |
| 30 | S4_R4 + LR | 5.283 | 5.099 | 0.921 | -0.521 |
| 31 | S0_R4 + RVRlin | 5.308 | 5.173 | 0.921 | -0.504 |
| 32 | S0_R4 + PCA + RVRlin | 5.309 | 5.172 | 0.921 | -0.504 |
| **33** | **S4_R8 + RR *** | **5.399** | **5.278** | **0.916** | **-0.511** |
| **34** | **S8_R4 + KRR *** | **5.435** | **5.252** | **0.913** | **-0.387** |
| 35 | S4_R8 + PCA + RR | 5.438 | 5.318 | 0.916 | -0.523 |
| 36 | S0_R4 + ENR | 5.446 | 5.301 | 0.914 | -0.528 |
| 37 | S0_R4 + PCA + RR | 5.447 | 5.268 | 0.916 | -0.549 |
| 38 | S8_R8 + PCA + KRR | 5.45 | 5.296 | 0.911 | -0.391 |

| | | | | | |
|---|---|---|---|---|---|
| 39 | S0_R4 + RR | 5.452 | 5.275 | 0.916 | -0.553 |
| 40 | S8_R4 + PCA + KRR | 5.459 | 5.318 | 0.916 | -0.397 |
| **41** | **S8_R8 + KRR *** | **5.462** | **5.296** | **0.913** | **-0.395** |
| **42** | **S0_R4 + LR *** | **5.483** | **5.327** | **0.914** | **-0.528** |
| 43 | S8_R4 + RVRpoly | 5.487 | 5.404 | 0.908 | -0.383 |
| 44 | S8_R8 + PCA + RVRpoly | 5.522 | 5.349 | 0.913 | -0.356 |
| 45 | S8_R8 + RVRpoly | 5.527 | 5.432 | 0.908 | -0.383 |
| 46 | S8_R8 + RVRlin | 5.535 | 5.39 | 0.908 | -0.378 |
| 47 | S8_R8 + PCA + RVRlin | 5.535 | 5.39 | 0.908 | -0.378 |
| 48 | S8_R4 + RVRlin | 5.542 | 5.38 | 0.911 | -0.391 |
| **49** | **S8_R4 + PCA + RVRlin *** | **5.543** | **5.375** | **0.909** | **-0.393** |
| **50** | **S8_R4 + PCA + RVRpoly *** | **5.546** | **5.451** | **0.909** | **-0.329** |
| 51 | S8_R4 + ENR | 5.555 | 5.333 | 0.914 | -0.53 |
| 52 | S8_R4 + LR | 5.574 | 5.336 | 0.914 | -0.523 |
| 53 | S0_R8 + PCA + GPR | 5.598 | 5.532 | 0.907 | -0.243 |
| 54 | S8_R4 + PCA + RR | 5.601 | 5.456 | 0.908 | -0.518 |
| 55 | S8_R4 + RR | 5.609 | 5.46 | 0.908 | -0.518 |
| 56 | S4_R8 + ENR | 5.631 | 5.433 | 0.911 | -0.53 |
| **57** | **S8_R8 + RR *** | **5.633** | **5.49** | **0.908** | **-0.526** |
| **58** | **S0_R8 + RVRpoly *** | **5.633** | **5.589** | **0.904** | **-0.453** |
| 59 | S0_R8 + GPR | 5.634 | 5.522 | 0.907 | -0.562 |
| 60 | S0_R8 + KRR | 5.634 | 5.585 | 0.904 | -0.477 |
| 61 | S0_R8 + RVRlin | 5.639 | 5.577 | 0.902 | -0.471 |
| 62 | S0_R8 + PCA + RVRlin | 5.639 | 5.577 | 0.902 | -0.471 |
| 63 | S0_R8 + PCA + KRR | 5.64 | 5.592 | 0.904 | -0.462 |
| 64 | S8_R8 + PCA + RR | 5.649 | 5.514 | 0.908 | -0.535 |
| **65** | **S0_R8 + PCA + RVRpoly *** | **5.663** | **5.607** | **0.903** | **-0.438** |
| **66** | **S4_R8 + LR *** | **5.664** | **5.466** | **0.911** | **-0.53** |
| 67 | S8_R8 + ENR | 5.695 | 5.467 | 0.906 | -0.533 |
| 68 | S8_R8 + LR | 5.716 | 5.466 | 0.908 | -0.528 |
| 69 | S0_R8 + RR | 5.806 | 5.707 | 0.902 | -0.564 |
| 70 | S0_R8 + PCA + RR | 5.833 | 5.732 | 0.902 | -0.569 |
| 71 | S4_R4 + PCA + ENR | 6.002 | 5.806 | 0.898 | -0.54 |
| 72 | S4_R4 + PCA + LR | 6.003 | 5.804 | 0.898 | -0.54 |
| **73** | **1273 + GPR *** | **6.052** | **5.977** | **0.887** | **-0.486** |
| **74** | **873 + GPR *** | **6.07** | **5.936** | **0.889** | **-0.476** |
| 75 | S0_R8 + ENR | 6.087 | 5.965 | 0.89 | -0.587 |
| 76 | 473 + GPR | 6.097 | 5.987 | 0.884 | -0.459 |
| 77 | S4_R8 + PCA + ENR | 6.119 | 5.88 | 0.893 | -0.546 |
| 78 | S0_R8 + LR | 6.121 | 5.996 | 0.89 | -0.581 |
| 79 | S4_R8 + PCA + LR | 6.122 | 5.878 | 0.893 | -0.546 |
| 80 | S8_R4 + PCA + ENR | 6.145 | 5.935 | 0.891 | -0.535 |
| **81** | **S8_R8 + PCA + ENR *** | **6.146** | **5.906** | **0.893** | **-0.533** |
| **82** | **S8_R4 + PCA + LR *** | **6.147** | **5.937** | **0.891** | **-0.535** |
| 83 | S8_R8 + PCA + LR | 6.15 | 5.917 | 0.893 | -0.533 |
| 84 | 473 + KRR | 6.175 | 6.107 | 0.883 | -0.423 |
| 85 | 173 + RVRpoly | 6.185 | 6.094 | 0.884 | -0.418 |

| | | | | | |
|---|---|---|---|---|---|
| 86 | 173 + KRR | 6.194 | 6.114 | 0.881 | -0.437 |
| 87 | 1273 + KRR | 6.196 | 6.157 | 0.882 | -0.447 |
| 88 | 173 + RVRlin | 6.21 | 6.116 | 0.882 | -0.418 |
| **89** | **473 + RVRpoly *** | **6.232** | **6.189** | **0.877** | **-0.418** |
| **90** | **1273 + RVRpoly *** | **6.237** | **6.186** | **0.88** | **-0.439** |
| 91 | 873 + KRR | 6.255 | 6.106 | 0.881 | -0.43 |
| 92 | 473 + ENR | 6.267 | 6.142 | 0.882 | -0.57 |
| 93 | 173 + RR | 6.277 | 6.187 | 0.881 | -0.593 |
| 94 | 1273 + RVRlin | 6.278 | 6.263 | 0.878 | -0.421 |
| 95 | 473 + RVRlin | 6.286 | 6.152 | 0.88 | -0.401 |
| 96 | 473 + RR | 6.288 | 6.201 | 0.881 | -0.583 |
| **97** | **473 + LR *** | **6.294** | **6.146** | **0.882** | **-0.563** |
| **98** | **873 + ENR *** | **6.299** | **6.195** | **0.882** | **-0.593** |
| 99 | 873 + RVRpoly | 6.31 | 6.195 | 0.882 | -0.433 |
| 100 | 1273 + ENR | 6.312 | 6.122 | 0.884 | -0.583 |
| 101 | 873 + LR | 6.316 | 6.226 | 0.882 | -0.588 |
| 102 | S4_R4 + RFR | 6.32 | 6.19 | 0.896 | -0.731 |
| 103 | 173 + ENR | 6.322 | 6.21 | 0.877 | -0.566 |
| 104 | 1273 + LR | 6.326 | 6.134 | 0.884 | -0.578 |
| **105** | **173 + GPR *** | **6.336** | **6.207** | **0.875** | **-0.466** |
| **106** | **173 + LR *** | **6.35** | **6.229** | **0.877** | **-0.556** |
| 107 | 873 + RVRlin | 6.366 | 6.272 | 0.877 | -0.413 |
| 108 | 1273 + RR | 6.372 | 6.27 | 0.877 | -0.593 |
| 109 | 873 + RR | 6.406 | 6.281 | 0.879 | -0.593 |
| 110 | S0_R4 + PCA + ENR | 6.478 | 6.383 | 0.881 | -0.623 |
| 111 | S0_R4 + PCA + LR | 6.488 | 6.384 | 0.881 | -0.623 |
| 112 | S8_R4 + RFR | 6.608 | 6.437 | 0.879 | -0.713 |
| **113** | **S0_R8 + PCA + ENR *** | **6.656** | **6.461** | **0.874** | **-0.618** |
| **114** | **S0_R8 + PCA + LR *** | **6.667** | **6.47** | **0.874** | **-0.618** |
| 115 | S4_R8 + RFR | 6.681 | 6.532 | 0.882 | -0.748 |
| 116 | S0_R4 + RFR | 6.805 | 6.695 | 0.884 | -0.77 |
| 117 | S8_R8 + RFR | 6.81 | 6.652 | 0.872 | -0.723 |
| 118 | S0_R4 + PCA + RFR | 7.306 | 7.11 | 0.85 | -0.742 |
| 119 | S0_R8 + RFR | 7.41 | 7.278 | 0.863 | -0.792 |
| 120 | S0_R8 + PCA + RFR | 7.552 | 7.354 | 0.843 | -0.773 |
| **121** | **S4_R4 + PCA + RFR *** | **7.662** | **7.464** | **0.833** | **-0.772** |
| **122** | **173 + RFR *** | **7.76** | **7.622** | **0.812** | **-0.733** |
| 123 | 1273 + RFR | 7.778 | 7.658 | 0.812 | -0.741 |
| 124 | S4_R8 + PCA + RFR | 7.802 | 7.624 | 0.833 | -0.796 |
| 125 | 873 + RFR | 7.802 | 7.654 | 0.815 | -0.74 |
| 126 | 473 + RFR | 7.84 | 7.693 | 0.815 | -0.743 |
| 127 | S8_R4 + PCA + RFR | 8.454 | 8.319 | 0.807 | -0.829 |
| 128 | S8_R8 + PCA + RFR | 8.477 | 8.378 | 0.809 | -0.834 |

**Table S3.** Cross-site results. Averaged CV MAE, test MAE and test correlation between predicted and true age for 32 selected workflows arranged in increasing order of test MAE (averaged over four datasets/runs). The 10 selected workflows for assessing test-retest reliability and longitudinal consistency are marked in bold letters with asterisk (*) .

| S.No. | Workflow Names | Average CV MAE | Average Test MAE | Averaged test correlation | Average test age-bias |
|---|---|---|---|---|---|
| **1** | **S4_R4 + PCA + GPR *** | **4.323** | **5.234** | **0.935** | **-0.269** |
| **2** | **S4_R4 + GPR *** | **4.275** | **5.357** | **0.932** | **-0.42** |
| **3** | **S4_R4 + PCA + RVRlin *** | **4.546** | **5.44** | **0.932** | **-0.378** |
| **4** | **S4_R4 + RVRlin *** | **4.544** | **5.464** | **0.932** | **-0.381** |
| **5** | **S4_R8 + RVRlin *** | **4.848** | **5.48** | **0.92** | **-0.362** |
| **6** | **S4_R4 + RR *** | **4.475** | **5.506** | **0.932** | **-0.439** |
| **7** | **S4_R4 + PCA + RR *** | **4.49** | **5.549** | **0.932** | **-0.449** |
| **8** | **S0_R4 + LR *** | **4.65** | **5.634** | **0.931** | **-0.452** |
| **9** | **S4_R8 + LR *** | **4.922** | **5.687** | **0.925** | **-0.467** |
| **10** | **S4_R8 + RR *** | **4.817** | **5.715** | **0.925** | **-0.473** |
| 11 | S8_R4 + PCA + GPR | 4.709 | 5.757 | 0.923 | -0.367 |
| 12 | S8_R4 + KRR | 4.79 | 5.795 | 0.92 | -0.393 |
| 13 | S8_R4 + PCA + RVRlin | 4.992 | 5.827 | 0.914 | -0.351 |
| 14 | S8_R4 + PCA + RVRpoly | 4.953 | 5.836 | 0.915 | -0.35 |
| 15 | S8_R8 + KRR | 4.839 | 5.872 | 0.92 | -0.4 |
| 16 | S0_R8 + PCA + RVRpoly | 5.236 | 5.956 | 0.908 | -0.377 |
| 17 | S0_R8 + RVRpoly | 5.211 | 5.988 | 0.911 | -0.377 |
| 18 | S8_R8 + RR | 4.969 | 6.134 | 0.917 | -0.467 |
| 19 | 1273 + GPR | 5.567 | 6.18 | 0.9 | -0.431 |
| 20 | 473 + RVRpoly | 5.693 | 6.252 | 0.896 | -0.423 |
| 21 | 873 + GPR | 5.586 | 6.261 | 0.896 | -0.419 |
| 22 | 473 + LR | 5.771 | 6.286 | 0.894 | -0.492 |
| 23 | S8_R8 + PCA + ENR | 5.269 | 6.347 | 0.91 | -0.448 |
| 24 | 1273 + RVRpoly | 5.758 | 6.366 | 0.895 | -0.409 |
| 25 | 873 + ENR | 5.718 | 6.378 | 0.894 | -0.507 |
| 26 | S8_R4 + PCA + LR | 5.229 | 6.47 | 0.911 | -0.44 |
| 27 | S0_R8 + PCA + LR | 5.615 | 6.522 | 0.898 | -0.509 |
| 28 | S0_R8 + PCA + ENR | 5.613 | 6.526 | 0.898 | -0.509 |
| 29 | 173 + LR | 6.025 | 6.816 | 0.883 | -0.497 |
| 30 | 173 + GPR | 6.153 | 6.978 | 0.866 | -0.474 |
| 31 | S4_R4 + PCA + RFR | 7.037 | 8.578 | 0.847 | -0.751 |
| 32 | 173 + RFR | 7.392 | 8.983 | 0.825 | -0.704 |

**Table S4.** Pearson correlation between true age difference and predicted age difference from two sessions at different test-retest durations and their respective mean absolute error (MAE) between true and predicted age for CoRR and OASIS-3 datasets for top 10 workflows.

| | **CoRR dataset** | | | **OASIS-3 dataset** | | |
|---|---|---|---|---|---|---|
| **Retest duration** | 2 – 3.25 years (N = 26) | | | 3 - 4 years (N = 127) | | |
| **Age range (years)** | 18.0 - 57.0 | | | 46.04 - 86.21 | | |
| **Workflows** | **MAE ses-1** | **MAE ses-2** | **correlation** | **MAE ses-1** | **MAE ses-2** | **correlation** |
| *S4_R4 + PCA + GPR* | *6.132* | *6.642* | *r = 0.447, p = 0.022* | *5.081* | *5.862* | *r = -0.008, p = 0.932* |
| S4_R4 + GPR | 6.539 | 7.044 | r = 0.444, p = 0.023 | 5.161 | 6.055 | r = -0.005, p = 0.955 |
| S4_R4 + PCA + RVRlin | 6.238 | 7.206 | r = 0.442, p = 0.024 | 5.869 | 6.928 | r = 0.063, p = 0.479 |
| S4_R4 + RVRlin | 6.229 | 7.201 | r = 0.451, p = 0.021 | 5.853 | 6.908 | r = 0.060, p = 0.499 |
| S4_R8 + RVRlin | 7.173 | 7.496 | r = 0.343, p = 0.086 | 5.669 | 6.962 | r = 0.027, p = 0.766 |
| S4_R4 + RR | 6.573 | 7.48 | r = 0.442, p = 0.024 | 5.634 | 6.601 | r = 0.051, p = 0.566 |
| S4_R4 + PCA + RR | 6.627 | 7.565 | r = 0.437, p = 0.026 | 5.735 | 6.673 | r = 0.051, p = 0.565 |
| S0_R4 + LR | 7.003 | 7.511 | r = 0.290, p = 0.151 | 5.81 | 6.726 | r = 0.006, p = 0.944 |
| S4_R8 + LR | 7.553 | 8.171 | r = 0.137, p = 0.505 | 6.571 | 7.572 | r = 0.095, p = 0.290 |
| S4_R8 + RR | 7.254 | 8.113 | r = 0.341, p = 0.088 | 5.999 | 6.871 | r = 0.058, p = 0.515 |

**Table S5**. Correlation of brain-age delta with various behavioral measures with and without bias correction. a. From within-site predictions. b. From cross-site predictions. Age, $age^2$, and gender were used as covariates. Abbreviations: CWIT: Color-Word Interference Test, TMT: Trail Making Test, WASI-II: Wechsler Abbreviated Scale of Intelligence

a. From within-site predictions

| Dataset | Behavioral measure | N | Correlation with age | No bias correction | | After bias correction | |
|---|---|---|---|---|---|---|---|
| | | | | (a) No covariate | (b) With covariates | (c) No covariate | (d) With covariates |
| CamCAN | Fluid Intelligence (Cattel test) | 631 | r = -0.661, p = 1.92e-80 | r = -0.043, p = 0.282 | **r = -0.131, p = 0.001** | **r = -0.157, p = 7.24e-05** | **r = -0.131, p = 0.001** |
| | Motor Learning (Reaction time) | 302 | r = 0.544, p = 1.11e-24 | r = 0.089, p = 0.122 | **r = 0.149, p = 0.010** | **r = 0.186, p = 0.001** | **r = 0.148, p = 0.010** |
| eNKI | CWIT (Inhibition trial completion time) | 340 | r = 0.361, p = 6.50e-12 | r = 0.022, p = 0.683 | **r = 0.113, p = 0.037** | r = 0.094, p = 0.084 | **r = 0.114, p = 0.035** |
| | TMT (Number Letter Switching trial completion time) | 344 | r = 0.279, p = 1.45e-07 | r = -0.031, p = 0.564 | r = 0.035, p = 0.520 | r = 0.022, p = 0.690 | r = 0.035, p = 0.522 |
| | WASI-II matrix reasoning | 347 | r = -0.240, p = 6.03e-06 | r = 0.026, p = 0.627 | r = -0.029, p = 0.597 | r = -0.019, p = 0.728 | r = -0.028, p = 0.607 |
| | WASI-II similarities | 347 | r = 0.052, p = 0.332 | r = -0.033, p = 0.536 | r = -0.02, p = 0.716 | r = -0.023, p = 0.667 | r = -0.019, p = 0.730 |

b. From cross-site predictions

| Dataset | Behavioral measure | N | Correlation with age | No bias correction | | After bias correction | |
|---|---|---|---|---|---|---|---|
| | | | | (a) No covariate | (b) With covariates | (c) No covariate | (d) With covariates |
| CamCAN | Fluid Intelligence (Cattel test) | 631 | r = -0.661, p = 1.92e-80 | r = 0.071, p = 0.074 | r = -0.068, p = 0.088 | r = -0.053, p = 0.180 | r = -0.070, p = 0.088 |
| | Motor Learning (Reaction time) | 302 | r = 0.544, p = 1.11e-24 | r = -0.023, p = 0.689 | r = 0.095, p = 0.100 | r = 0.083, p = 0.151 | r = 0.095, p = 0.100 |
| eNKI | CWIT (Inhibition trial completion time) | 340 | r = 0.361, p = 6.50e-12 | r = 0.005, p = 0.931 | **r = 0.180, p = 0.001** | r = 0.065, p = 0.230 | **r = 0.180, p = 0.001** |
| | TMT (Number Letter Switching trial completion time) | 344 | r = 0.279, p = 1.45e-07 | r = -0.007, p = 0.898 | **r = 0.137, p = 0.011** | r = 0.039, p = 0.469 | **r = 0.137, p = 0.011** |
| | WASI-II matrix reasoning | 347 | r = -0.240, p = 6.03e-06 | r = 0.077, p = 0.150 | r = -0.048, p = 0.371 | r = 0.043, p = 0.425 | r = -0.048, p = 0.371 |
| | WASI-II similarities | 347 | r = 0.052, p = 0.332 | r = -0.098, p = 0.068 | r = -0.090, p = 0.096 | r = -0.096, p = 0.073 | r = -0.090, p = 0.096 |

**Table S6.** Pearson's correlation coefficients between corrected brain-age delta using S4_R4 + PCA + GPR workflow and cognitive measures using age, age$^2$ and gender as covariates from th the ADNI sample. The correlations with disease severity (MMSE and CDR) and ability to function independently (FAQ) were computed for the whole sample and each diagnostic group (CN, EMCI, LMCI and AD) separately from two time points. Abbreviations: MMSE: Mini-Mental State Examination, CDR: Global Clinical Dementia Rating Scale, FAQ: Functional Assessment Questionnaire; CN: cognitively normal, EMCI: early mild cognitive impairment, LMCI: late mild cognitive impairment, AD: Alzheimer's disease

| | **Timepoint 1 (T1)** | | | **Timepoint 2 (T2)** | | |
|---|---|---|---|---|---|---|
| | **MMSE** | **CDR** | **FAQ** | **MMSE** | **CDR** | **FAQ** |
| **CN** | N = 68 | N = 67 | N = 74 | N = 153 | N = 147 | N = 149 |
| | r = -0.170, p = 0.167 | r = -0.0149, p = 0.904 | r = 0.134, p = 0.255 | r = -0.064, p = 0.430 | r = -0.008, p = 0.925 | r = 0.073, p = 0.372 |
| **EMCI** | N = 3 | N = 3 | N = 3 | N = 196 | N = 194 | N = 193 |
| | n.a. | n.a. | n.a. | r = -0.102, p = 0.154 | **r = 0.147, p = 0.041** | r = 0.079, p = 0.277 |
| **LMCI** | N = 2 | N = 2 | N = 2 | N = 103 | N = 102 | N = 103 |
| | n.a. | n.a. | n.a. | **r = -0.258, p = 0.008** | r = 0.126, p = 0.207 | **r = 0.302, p = 0.002** |
| **AD** | N = 17 | N = 17 | N = 26 | N = 61 | N = 61 | N = 61 |
| | r = -0.401, p = 0.111 | r = 0.338, p = 0.184 | r = 0.247, p = 0.223 | r = -0.178, p = 0.169 | r = 0.202, p = 0.119 | **r = 0.303, p = 0.018** |
| **Whole sample** | N = 90 | N = 89 | N = 105 | N = 513 | N = 504 | N = 506 |
| | **r = -0.229, p = 0.030** | r = 0.100, p = 0.352 | **r = 0.277, p = 0.004** | **r = -0.319, p = 1.34e-13** | **r = 0.280, p = 1.65e-10** | **r = 0.338, p = 5.38e-15** |

**Figure S1.** Histogram of age of the data used for final model training.

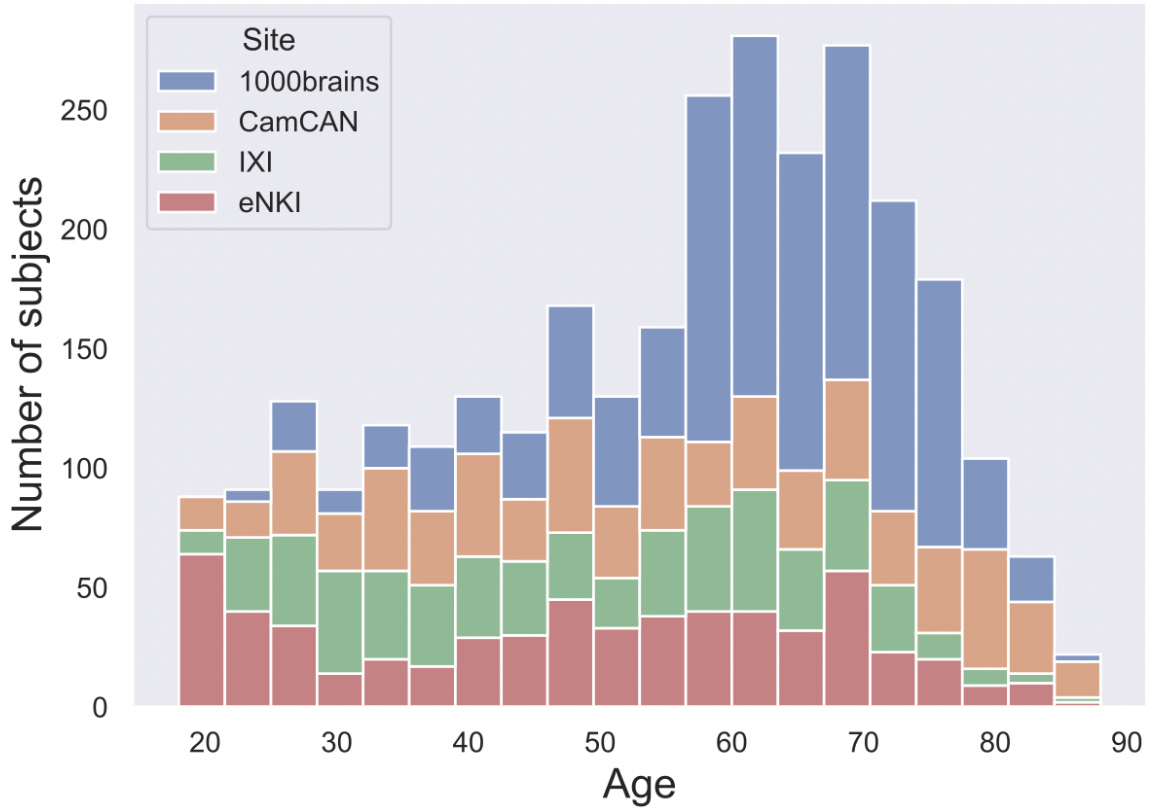

**Figure S2.** Correlation between predictions from different workflows using CamCAN data. a. 128 workflows from within-site analysis b. 32 workflows from cross-site analysis

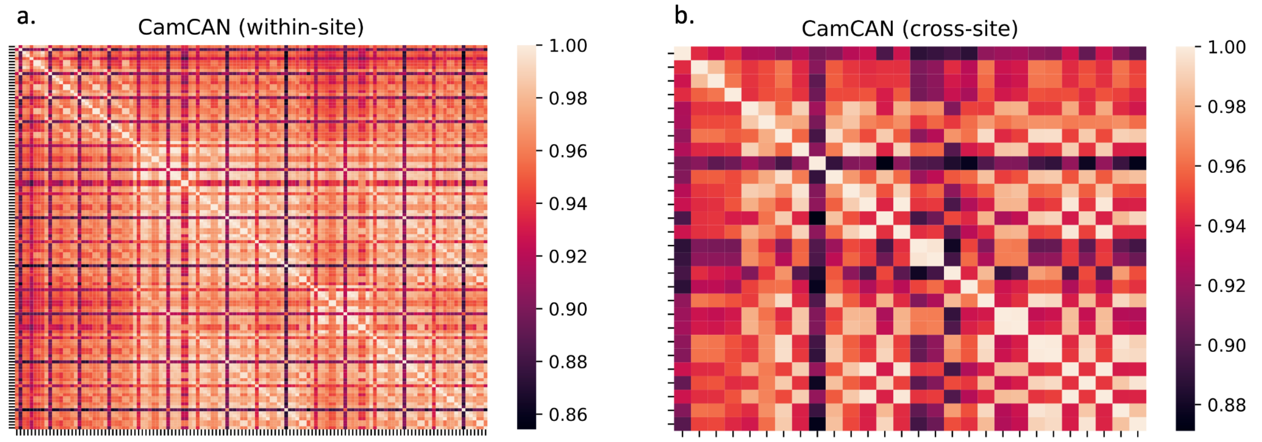


Labels for within-site workflows list as plotted in the heatmap above (a)

| | | | | | |
|---|---|---|---|---|---|
| 1 | 173 + RR | 44 | S0_R4 + PCA + KRR | 87 | S0_R8 + ENR |
| 2 | 173 + RFR | 45 | S0_R4 + PCA + GPR | 88 | S0_R8 + RVRpoly |
| 3 | 173 + RVRlin | 46 | S0_R4 + PCA + LR | 89 | S0_R8 + PCA + RR |
| 4 | 173 + KRR | 47 | S0_R4 + PCA + ENR | 90 | S0_R8 + PCA + RFR |
| 5 | 173 + GPR | 48 | S0_R4 + PCA + RVRpoly | 91 | S0_R8 + PCA + RVRlin |
| 6 | 173 + LR | 49 | S4_R4 + RR | 92 | S0_R8 + PCA + KRR |
| 7 | 173 + ENR | 50 | S4_R4 + RFR | 93 | S0_R8 + PCA + GPR |
| 8 | 173 + RVRpoly | 51 | S4_R4 + RVRlin | 94 | S0_R8 + PCA + LR |
| 9 | 473 + RR | 52 | S4_R4 + KRR | 95 | S0_R8 + PCA + ENR |
| 10 | 473 + RFR | 53 | S4_R4 + GPR | 96 | S0_R8 + PCA + RVRpoly |
| 11 | 473 + RVRlin | 54 | S4_R4 + LR | 97 | S4_R8 + RR |
| 12 | 473 + KRR | 55 | S4_R4 + ENR | 98 | S4_R8 + RFR |
| 13 | 473 + GPR | 56 | S4_R4 + RVRpoly | 99 | S4_R8 + RVRlin |
| 14 | 473 + LR | 57 | S4_R4 + PCA + RR | 100 | S4_R8 + KRR |
| 15 | 473 + ENR | 58 | S4_R4 + PCA + RFR | 101 | S4_R8 + GPR |
| 16 | 473 + RVRpoly | 59 | S4_R4 + PCA + RVRlin | 102 | S4_R8 + LR |
| 17 | 873 + RR | 60 | S4_R4 + PCA + KRR | 103 | S4_R8 + ENR |
| 18 | 873 + RFR | 61 | S4_R4 + PCA + GPR | 104 | S4_R8 + RVRpoly |
| 19 | 873 + RVRlin | 62 | S4_R4 + PCA + LR | 105 | S4_R8 + PCA + RR |
| 20 | 873 + KRR | 63 | S4_R4 + PCA + ENR | 106 | S4_R8 + PCA + RFR |
| 21 | 873 + GPR | 64 | S4_R4 + PCA + RVRpoly | 107 | S4_R8 + PCA + RVRlin |
| 22 | 873 + LR | 65 | S8_R4 + RR | 108 | S4_R8 + PCA + KRR |
| 23 | 873 + ENR | 66 | S8_R4 + RFR | 109 | S4_R8 + PCA + GPR |
| 24 | 873 + RVRpoly | 67 | S8_R4 + RVRlin | 110 | S4_R8 + PCA + LR |
| 25 | 1273 + RR | 68 | S8_R4 + KRR | 111 | S4_R8 + PCA + ENR |
| 26 | 1273 + RFR | 69 | S8_R4 + GPR | 112 | S4_R8 + PCA + RVRpoly |
| 27 | 1273 + RVRlin | 70 | S8_R4 + LR | 113 | S8_R8 + RR |
| 28 | 1273 + KRR | 71 | S8_R4 + ENR | 114 | S8_R8 + RFR |
| 29 | 1273 + GPR | 72 | S8_R4 + RVRpoly | 115 | S8_R8 + RVRlin |
| 30 | 1273 + LR | 73 | S8_R4 + PCA + RR | 116 | S8_R8 + KRR |
| 31 | 1273 + ENR | 74 | S8_R4 + PCA + RFR | 117 | S8_R8 + GPR |
| 32 | 1273 + RVRpoly | 75 | S8_R4 + PCA + RVRlin | 118 | S8_R8 + LR |
| 33 | S0_R4 + RR | 76 | S8_R4 + PCA + KRR | 119 | S8_R8 + ENR |
| 34 | S0_R4 + RFR | 77 | S8_R4 + PCA + GPR | 120 | S8_R8 + RVRpoly |
| 35 | S0_R4 + RVRlin | 78 | S8_R4 + PCA + LR | 121 | S8_R8 + PCA + RR |
| 36 | S0_R4 + KRR | 79 | S8_R4 + PCA + ENR | 122 | S8_R8 + PCA + RFR |
| 37 | S0_R4 + GPR | 80 | S8_R4 + PCA + RVRpoly | 123 | S8_R8 + PCA + RVRlin |
| 38 | S0_R4 + LR | 81 | S0_R8 + RR | 124 | S8_R8 + PCA + KRR |
| 39 | S0_R4 + ENR | 82 | S0_R8 + RFR | 125 | S8_R8 + PCA + GPR |
| 40 | S0_R4 + RVRpoly | 83 | S0_R8 + RVRlin | 126 | S8_R8 + PCA + LR |
| 41 | S0_R4 + PCA + RR | 84 | S0_R8 + KRR | 127 | S8_R8 + PCA + ENR |
| 42 | S0_R4 + PCA + RFR | 85 | S0_R8 + GPR | 128 | S8_R8 + PCA + RVRpoly |
| 43 | S0_R4 + PCA + RVRlin | 86 | S0_R8 + LR | | |

Labels for cross-site workflows list as plotted in the heatmap above (b)

| | | | | | |
|---|---|---|---|---|---|
| 1 | 173 + RFR | 12 | S4_R4 + RVRlin | 23 | S0_R8 + RVRpoly |
| 2 | 173 + GPR | 13 | S4_R4 + GPR | 24 | S0_R8 + PCA + LR |
| 3 | 173 + LR | 14 | S4_R4 + PCA + RR | 25 | S0_R8 + PCA + ENR |
| 4 | 473 + LR | 15 | S4_R4 + PCA + RFR | 26 | S0_R8 + PCA + RVRpoly |
| 5 | 473 + RVRpoly | 16 | S4_R4 + PCA + RVRlin | 27 | S4_R8 + RR |
| 6 | 873 + GPR | 17 | S4_R4 + PCA + GPR | 28 | S4_R8 + RVRlin |
| 7 | 873 + ENR | 18 | S8_R4 + KRR | 29 | S4_R8 + LR |
| 8 | 1273 + GPR | 19 | S8_R4 + PCA + RVRlin | 30 | S8_R8 + RR |
| 9 | 1273 + RVRpoly | 20 | S8_R4 + PCA + GPR | 31 | S8_R8 + KRR |
| 10 | S0_R4 + LR | 21 | S8_R4 + PCA + LR | 32 | S8_R8 + PCA + ENR |
| 11 | S4_R4 + RR | 22 | S8_R4 + PCA + RVRpoly | | |

**Figure S3.** Bias correction. *(top)* within-site: *(left)* scatter plot between brain-age delta and age, *(right)* scatter plot between corrected brain-age delta and age, *(bottom)* cross-site: *(left)* scatter plot between brain-age delta and age, *(right)* scatter plot between corrected brain-age delta and age. The workflow used here is S4_R4 + PCA + GPR. a. CamCAN dataset b. eNKI dataset

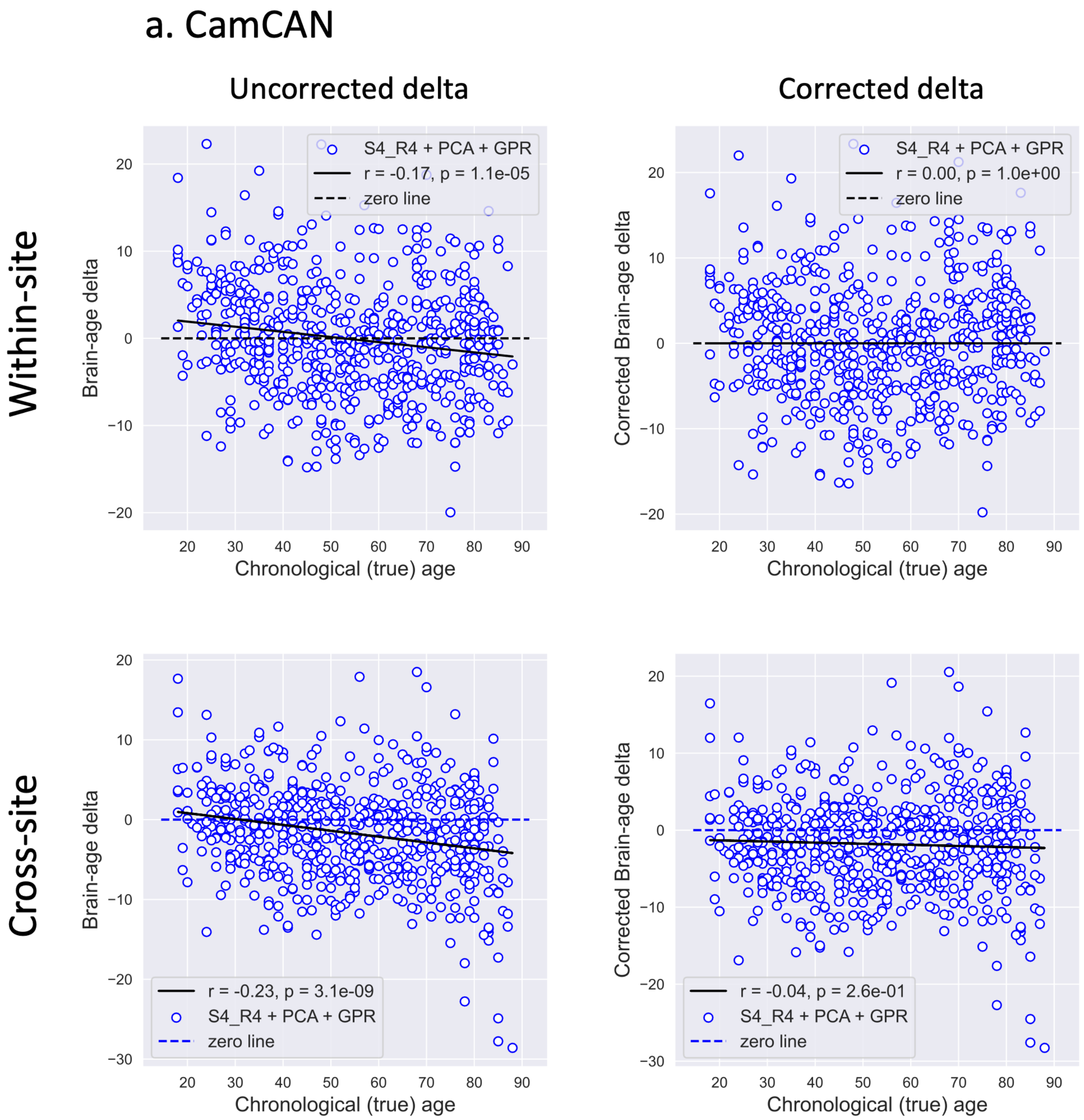

b. eNKI

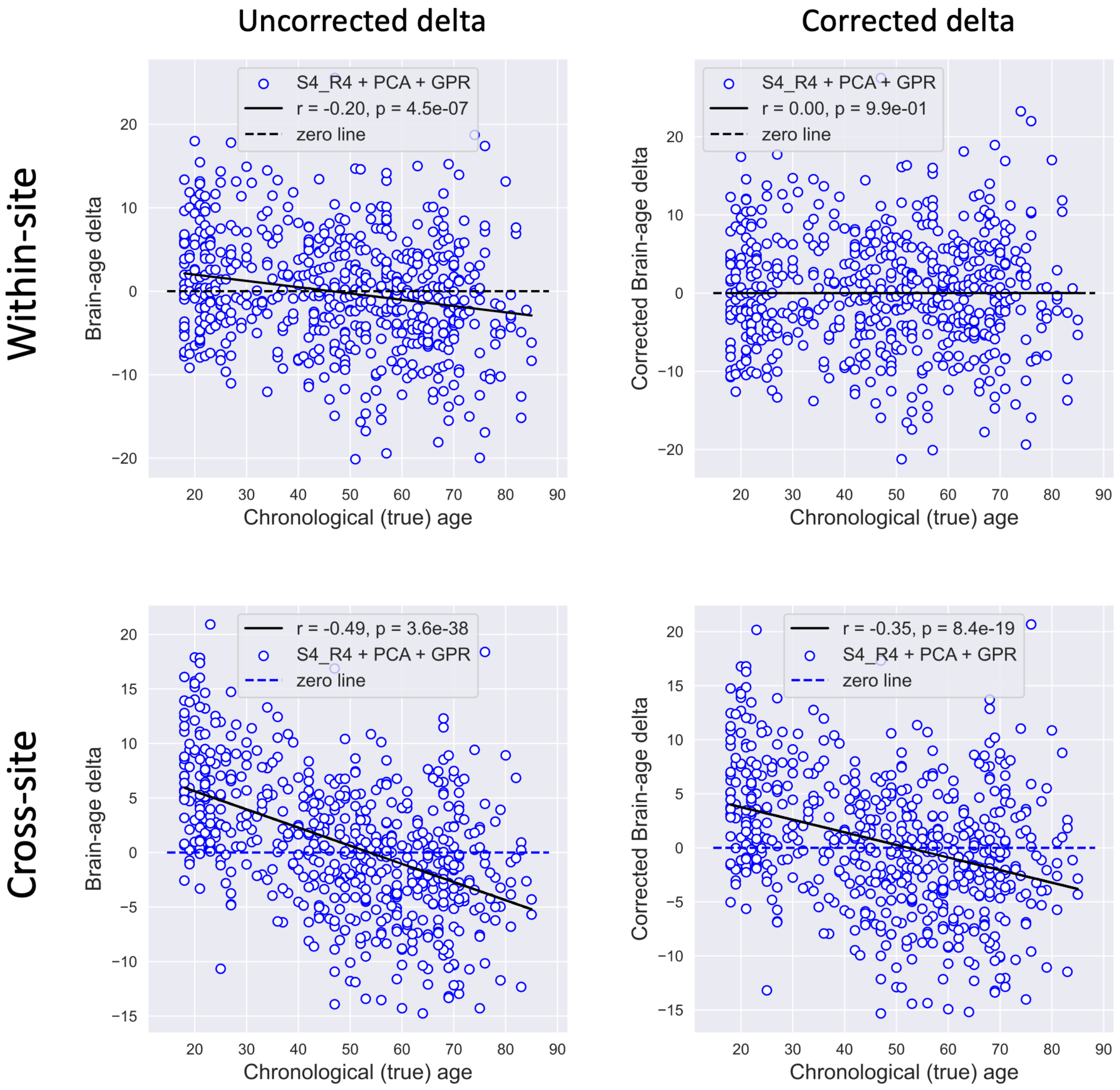

Uncorrected delta
Corrected delta
Within-site
Cross-site
S4_R4 + PCA + GPR
r = -0.20, p = 4.5e-07
zero line
r = 0.00, p = 9.9e-01
r = -0.49, p = 3.6e-38
r = -0.35, p = 8.4e-19
Brain-age delta
Corrected Brain-age delta
Chronological (true) age

**Figure S4.** CV performance of 128 workflows from within-site analysis. a. grouped using feature spaces b. grouped using ML algorithms

a.

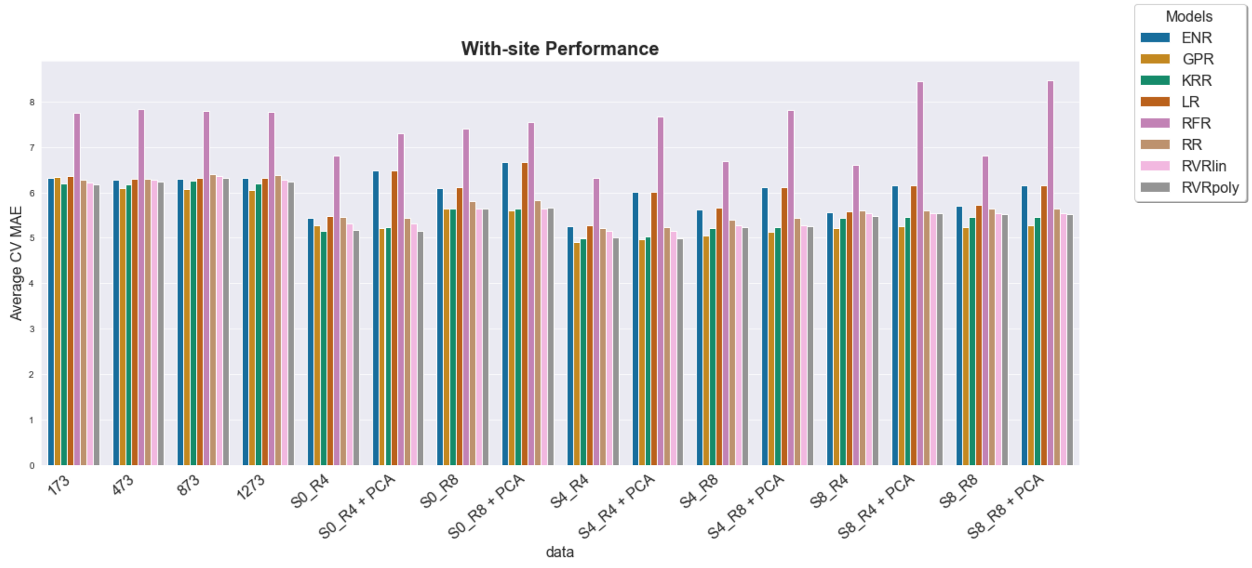


b.

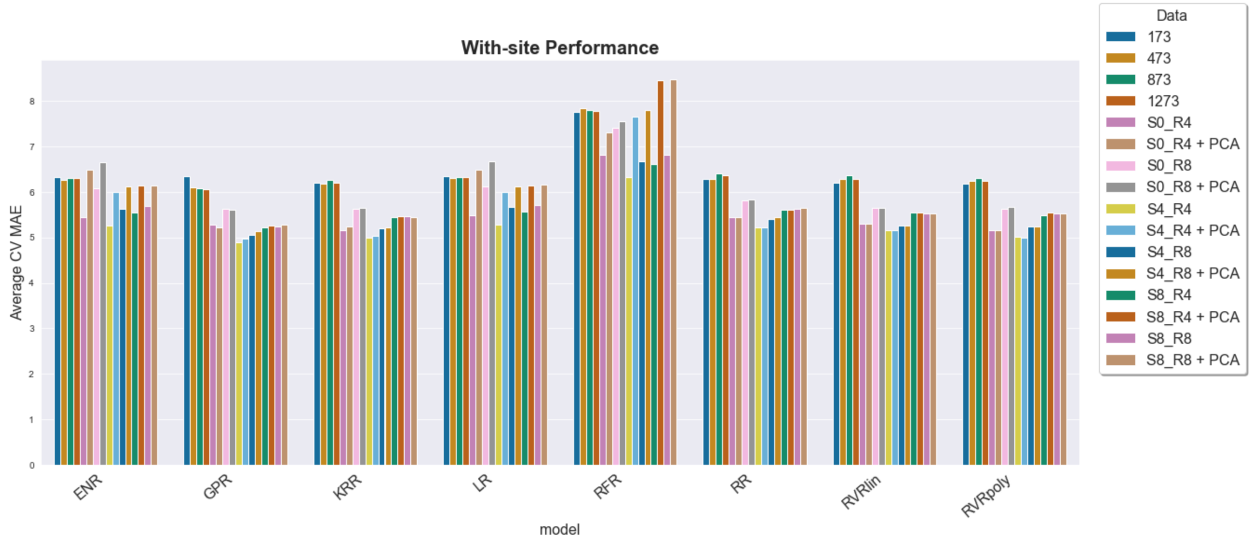

**Figure S5.** Correlation between MAE and age-bias (correlation between brain-age delta and true age) on CamCAN data. a. 128 workflows from within-site analysis b. 32 workflows from cross-site analysis

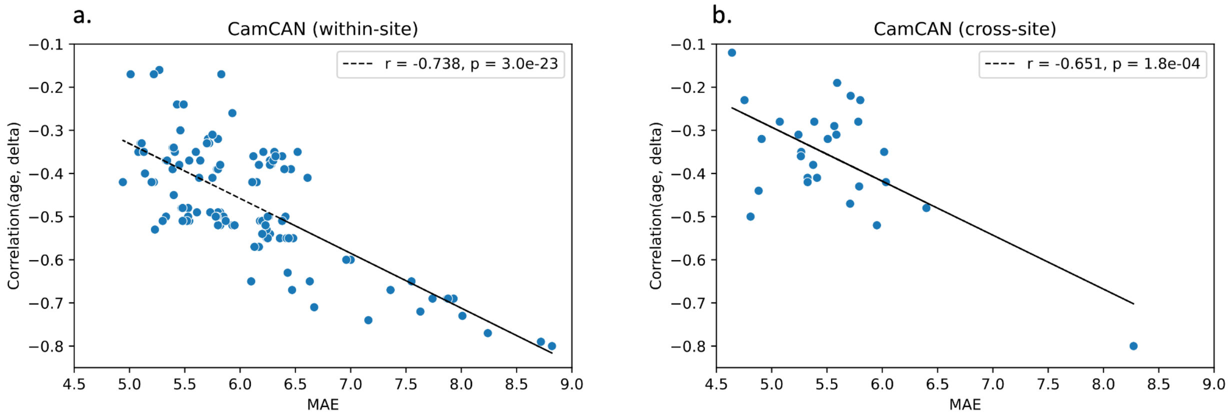

**Figure S6.** Correlation between within-site predictions and cross-site predictions using S4_R4 + PCA + GPR workflow a. CamCAN dataset b. eNKI dataset

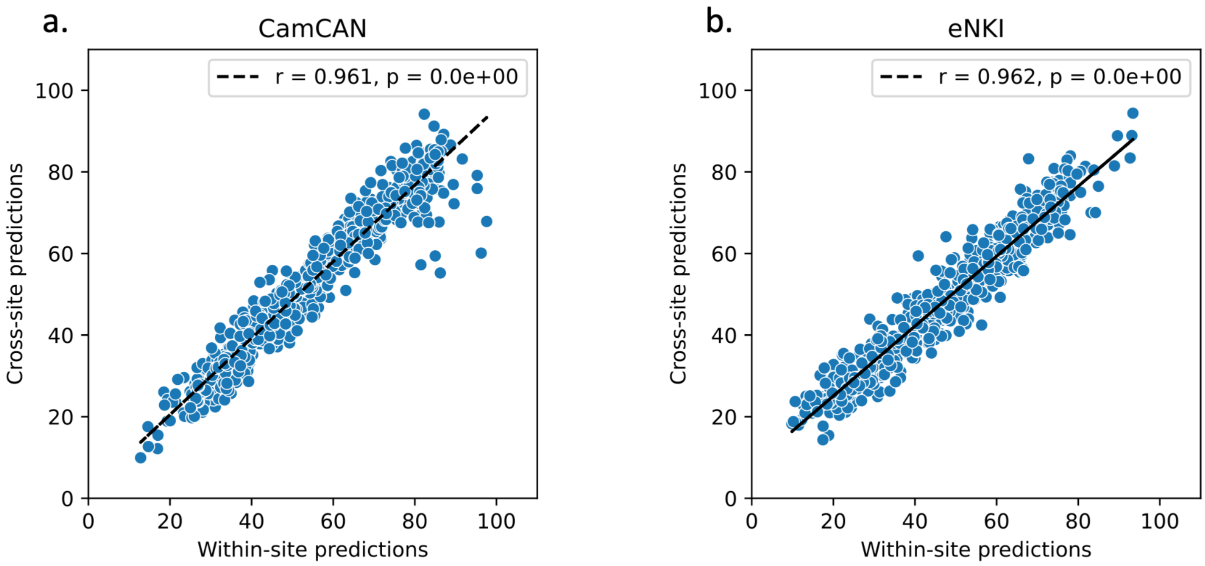

**Figure S7.** Effect of the number of samples used for bias correction. Different sub-sample of the healthy subjects (ADNI-CN) were drawn (0.1 to 0.9 fraction in steps of 0.1) without replacement, repeated 100 times. Bias correction parameters were computed using these CN subsamples and used to correct the bias in all ADNI-CN and ADNI-AD subjects. a. ADNI timepoint-T1: N for ADNI-CN for each sample fraction: 0.1 = 21, 0.2 = 42, 0.3 = 63, 0.4 = 84, 0.5 = 104, 0.6 = 125, 0.7 = 146, 0.8 = 167, 0.9 = 188, 1 = 209 and ADNI-AD N = 125, b. ADNI timepoint-T2: N for ADNI-CN for each sample fraction: 0.1 = 15, 0.2 = 31, 0.3 = 46, 0.4 = 61, 0.5 = 76, 0.6 = 92, 0.7 = 107, 0.8 = 122, 0.9 = 138, 1 = 153 and ADNI-AD N = 61. Each dot represents the mean brain-age delta of *(top)* all CN subjects and *(bottom)* all AD subjects (y-axis) when different numbers of samples from CN subjects (x-axis) are used for bias correction. CN: cognitively normal, AD: Alzheimer's Disease

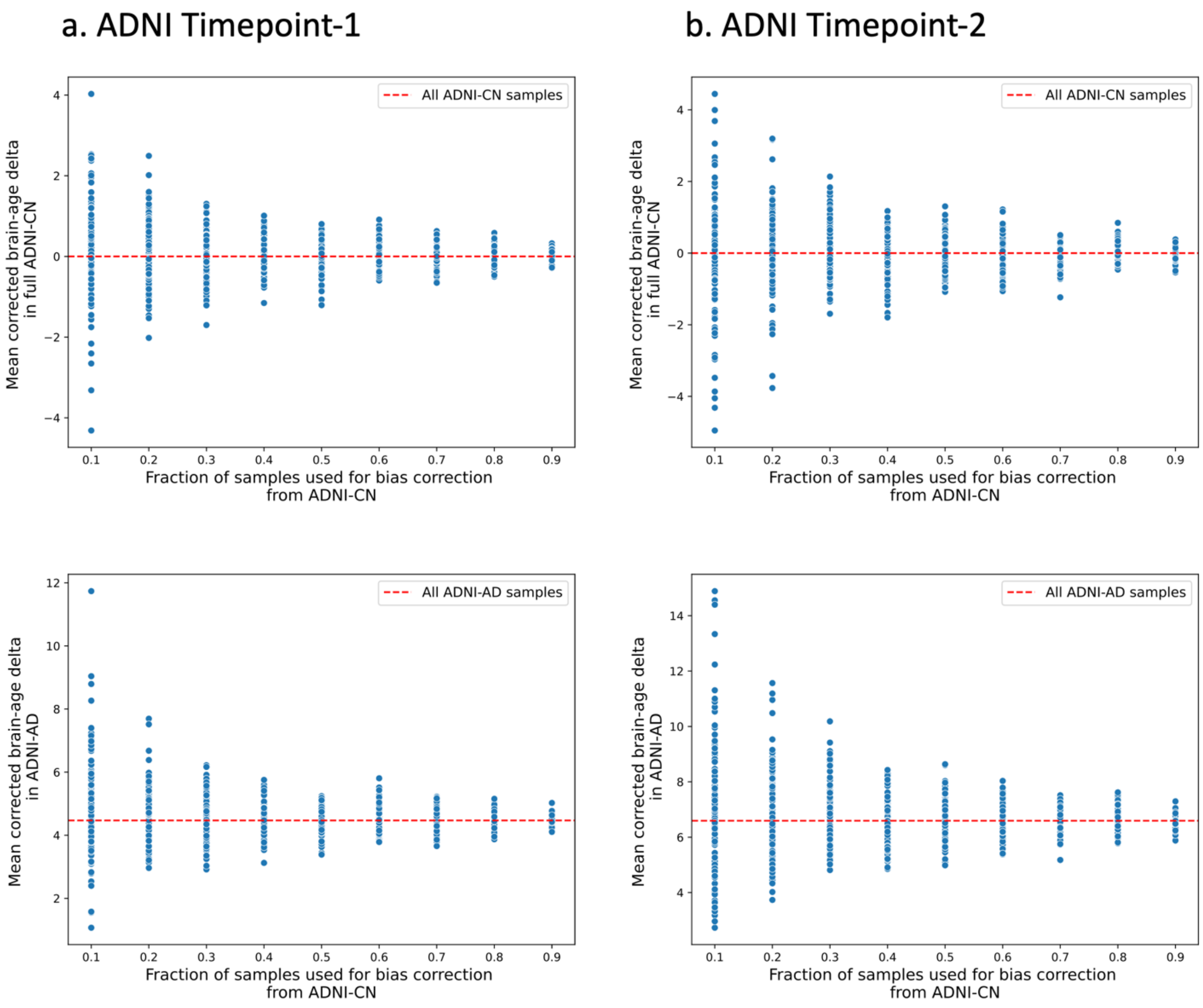

### Methods

#### 1. Machine Learning algorithms

Machine Learning (ML) algorithms used in the current study includes:

**1. Ridge regression (RR)** is a regularized linear model that minimizes the sum of the squared prediction error and employs an L2-norm regularization, i.e., minimizes the sum of the squares of regression coefficients (Hoerl and Kennard, 1970). The L2-norm helps to reduce the model complexity by shrinking some coefficients towards zero. The hyperparameter λ controls the regularization extent.

**2. LASSO regression (LR)** is another form of regularized linear regression that minimizes the sum of the squared prediction error and employs an L1-norm regularization i.e., minimizes the sum of the absolute values of the regression coefficients. The L1-norm regularization can drive some coefficients to zero, resulting in a sparse model that helps mitigate multicollinearity and reduce model complexity (Tibshirani, 2011). The extent of regularization is determined by the hyperparameter λ.

**3. Elastic net regression (ENR):** It uses a weighted combination of L1 and L2 regularization in the Ordinary Least Squares (OLS) loss function. The relative weighting of the L1-norm and L2-norm contributions is controlled by a hyperparameter $\alpha$ (between 0 and 1) (Zou and Hastie, 2005). This shrinks some coefficients and sets some to zero resulting in a sparse model. Another hyperparameter λ controls the weighting of the sum of both penalties to the loss function.

**4. Kernel ridge regression (KRR):** KRR combines ridge regression with the kernel trick (Murphy, 2012). It aims to learn a function in the space induced by the respective kernel by minimizing a squared loss with a squared norm regularization term. The kernel was set to polynomial. The kernel degree (1 or 2) and the level of L2 regularization $\alpha$ were the hyperparameters.

**5. Gaussian processes regression (GPR):** GPR incorporates nonlinear and Gaussian probabilistic elements to allow quantitative predictions to be made using continuous variables (Rasmussen, 2004; Doyle *et al.*, 2013). GPR infers a probability distribution of the target. It requires specifying a prior distribution on the parameter as a mean and covariance kernel and relocates probabilities based on the training data using the Bayes' rule. The prior distribution is usually assumed to be a multivariate Gaussian distribution with zero mean. We used a radial basis kernel, and the number of restarts was set to 100 for finding the kernel parameters (length scale and bound), maximizing the log-marginal likelihood.

**6. Relevance vector regression (RVR):** RVR is a Bayesian sparse learning model analogous to Support Vector Machine (Tipping, 2000). It assumes the prior probability distribution of the weights of the input data as a zero-mean normal distribution and iteratively adjusts the values of precision in the model using evidence approximation. The weights with weak precision are set to zero in the training and provide few basis functions. Since RVR has no algorithm-specific hyperparameters, the training process is computationally efficient. We used two types of kernel for RVR; linear kernel (RVRlin) and polynomial kernel of degree 1 (RVRpoly).

**7. Random forest regression (RFR):** Random forest (RF) is an ensemble technique that uses multiple regression trees trained with subsets of data generated with the bagging (Bootstrap aggregating) technique (Breiman, 2001). Predictions are made by averaging the predictions of each tree. RFR can capture non-linear or complex relationships between features and labels and is known to be resilient to overfitting.

#### 2. Behavioral/Cognitive measures

##### 2.1 From CamCAN dataset:

From the Cam-CAN dataset (Taylor *et al.*, 2017) we used two behavioral measures:

1. Fluid Intelligence (FI; N = 631) assessed by the accuracy on the Cattell Culture Fair Test, which contains four subtests of nonverbal puzzles involving series completion, classification, matrices, and conditions. One participant scoring below 12 was deemed non-engaging and removed from the analysis.
2. Reaction time for the motor learning task (N = 302), which was pressured movement of a cursor to a target by moving an (occluded) stylus under veridical, perturbed (30°), and reset (veridical again) mappings between visual and real space.

##### 2.2 From eNKI dataset:

From the eNKI dataset, we used measures for cognitive and executive functioning assessed by various tests from the Delis-Kaplan Executive Functioning System (D-KEFS), and cognitive intelligence measured by Wechsler Abbreviated Scale of Intelligence (WASI-II) (Nooner *et al.*, 2012). We included two tests from the D-KEFS battery: Colour-Word Interference Test (CWIT) and Trail Making Test (TMT). The WASI-II consists of four subtests: vocabulary, similarities, block design, and matrix reasoning. Specifically, we correlated brain-age delta with:

1. The completion time of CWIT inhibition trial (N=340), which measures inhibitory control and selective attention.
2. The completion time of the D-KEFS TMT number-letter switching condition (N=344), which measures cognitive flexibility.
3. WASI-II similarities (N=347), which is one of the subcomponents of crystallized intelligence.
4. WASI-II matrix reasoning (N=347), which is one of the subcomponents of nonverbal fluid abilities and visuomotor/coordination skills.
